# State-space trajectories and traveling waves following distraction

**DOI:** 10.1101/2024.02.19.581020

**Authors:** Tamal Batabyal, Scott L. Brincat, Jacob A. Donoghue, Mikael Lundqvist, Meredith K. Mahnke, Earl K. Miller

## Abstract

Cortical activity shows the ability to recover from distractions. We analyzed neural activity from the prefrontal cortex (PFC) of monkeys performing working memory tasks with mid-memory-delay distractions (a cued gaze shift or an irrelevant visual input). After distraction there were state-space rotational dynamics that returned spiking to population patterns similar to those pre-disruption. In fact, rotations were fuller when the task was performed correctly versus when errors were made. We found a correspondence between state-space rotations and traveling waves across the surface of the PFC. This suggests a role for emergent dynamics like state-space rotations and traveling waves in recovery from distractions.

## Introduction

The cortex manages to maintain consistent neural coding in the face of external perturbations. Neural activity converges to a sequence of states that produce consistent computations despite noise, variation, and disruption (Kozachkov et al., 2020; Spaak et al., 2017; Stokes, 2015; Stokes et al., 2013; Tabareau et al., 2010; Wasmuht et al., 2018). This consistency is reflected in subspace coding. The high-dimensional spiking of many neurons can be reduced to just a few meaningful dimensions. This indicates a great deal of coordination of spiking across neurons at slower time scales (Börgers & Kopell, 2003; Denève & Machens, 2016; Eisen et al., 2024; Kozachkov et al., 2020; Mariño et al., 2005; Ruan et al., 2014; Shu et al., 2003; Sussillo, 2014; Turrigiano & Nelson, 2004). Across much of the cortex, these state-space dynamics show rotational components (Aoi et al., 2020; Churchland et al., 2012; Gao et al., 2016; Hall et al., 2014; Jiang et al., 2020; Libby & Buschman, 2021; Mante et al., 2013). This indicates that spiking activity forms orderly, sequential patterns. When the rotations form a complete cycle, population activity returns to, or near to, a previous state.

We used working memory for a test of whether recovery from distractions might involve such rotational dynamics. Memory maintenance can be disrupted by distractions, yet neural activity often, but not always, manages to recover and return to a pre-disruption state that restores focus on the task (Derrfuss et al., 2017; Lewis-Peacock et al., 2015; Mallett & Lewis-Peacock, 2019; Parthasarathy et al., 2017; Suzuki & Gottlieb, 2013; Yoon et al., 2006). We examined activity from the prefrontal cortex of non-human primates (NHPs; Rhesus Macaque) performing two working memory tasks with a mid-memory delay distraction: a saccade or a visual distractor. We found that population spiking activity showed rotational dynamics that were predictive of behavioral performance. We also found a correspondence between rotations in population state-space and traveling waves propagating across the cortical surface.

## Results

### Rotational dynamics after a saccade distraction

We first examined a visual working memory task that included (on half of randomly selected trials) a cued shift in gaze in the middle of the memory delay (Fig. 1a (Brincat et al., 2021)). Neural activity was recorded from the dorsolateral and ventrolateral PFC (dlPFC and vlPFC) and from both hemispheres simultaneously. The trial began with fixation on a point to the left or right (50% of trials randomly) of a computer screen. An object briefly appeared as a sample in the center of the screen, thus in the right or left visual hemifield, respectively. The sample could be one of two different objects, at one of two different locations slightly above or below the center of the screen. Subjects were required to remember both the object identity and location over a blank 1.6 s working memory delay. Thus, there were four working memory conditions— two object identities ⨉ two object locations (upper vs. lower). On 50% of the trials, at the midpoint of the delay period, the fixation point jumped across the vertical midline to the opposite side, cueing a gaze shift to its new location. This shifted the remembered location of the sample object from one visual hemifield to the other. At the end of the delay, a test object appeared.

**Figure 1.**
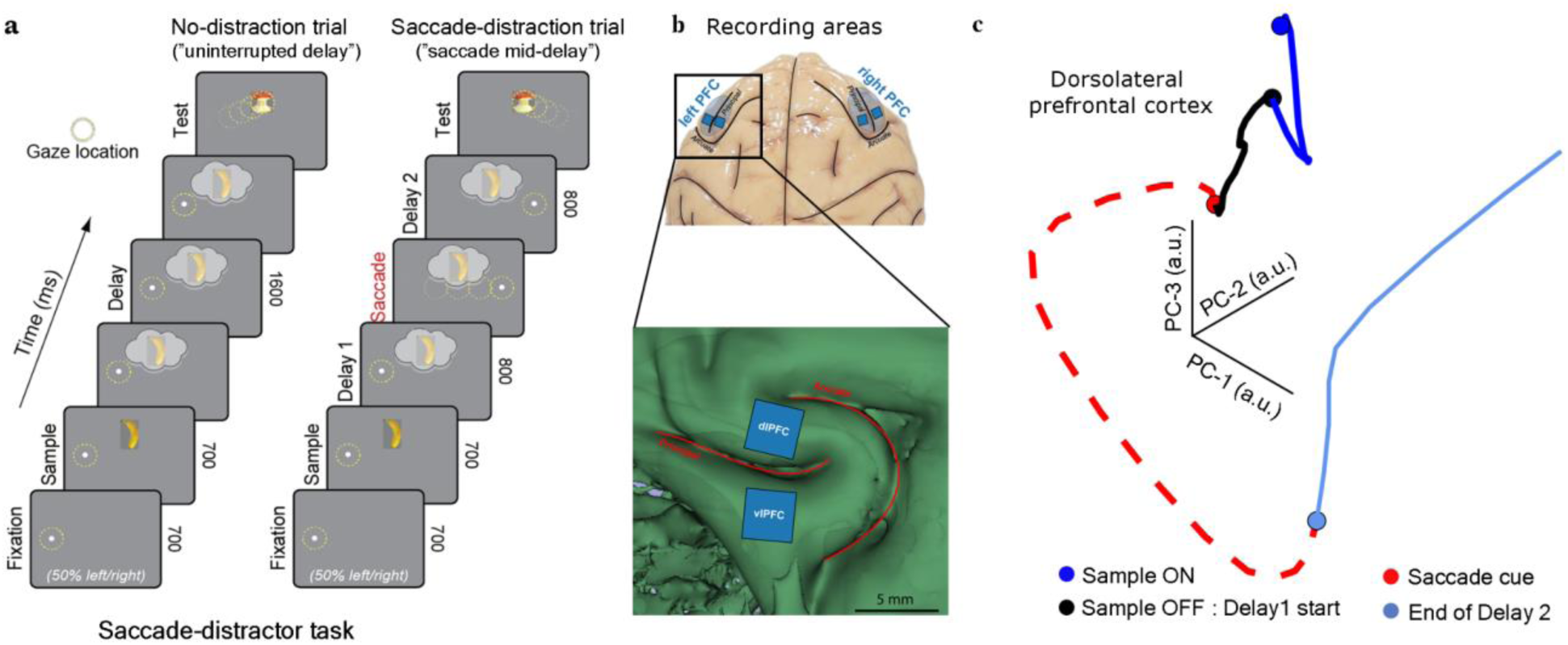
The saccade-distractor task and rotational dynamics. (a) A working memory task with a saccade in the middle of the delay. Subjects fixated to the left or the right of the center of the screen while a sample stimulus (randomly picked from one two images, Obj1 or Obj2) was shown above or below the screen center. On half of the randomly-chosen trials a saccade instructed shift of gaze to the opposite side of the screen. We focus on spiking after this saccade (“Delay 2”). (b) (*upper*) Schematic of recording locations, showing the left and right PFC areas. The arrays were marked by blue squares. (*lower)* magnified lateral PFC region with scale bar. (c) Population spiking activity across the duration of the trial, projected onto its top 3 principal components, for an example session.

Subjects were required to saccade to it if it did not match the remembered sample in either object identity or upper/lower location, and otherwise withhold response until a subsequently presented non-matching test object.

We previously found evidence that the cued gaze shift transferred the neural representation of working memories from the corresponding (contralateral to the sample) “sending” hemisphere to the other “receiving” hemisphere (Brincat et al., 2021). But evidence suggests the mid-delay saccade also acted as a distractor that impaired working memory. Performance was significantly poorer on trials with the saccade (accuracy = 80.33 ± 0.06%, mean ± s.e.m.) than trials without it (90.10 ± 0.04%)(Brincat et al., 2021). Here, our main interest was how cortical spiking activity recovered from this distraction. Thus, our analyses focus on trials with a cued saccade and the 800 ms from the cue to the end of the memory delay (“Delay 2” in Fig. 1a). We also first focus on correctly performed trials.

To visualize the neural dynamics induced by the distracting saccade, we first projected population spiking activity onto its top three principal components (PCs) across time for one example recording session (Fig. 1b). The resulting neural trajectories suggested the distractor induced rotational dynamics in the neural population activity. After the saccade (Delay 2, dashed line in Fig. 1b), neural trajectories follow a smooth, elongated curve. This population activity motif is known as a rotation (Churchland et al., 2012). Such rotations reflect an orderly progression of activity through the neural population and a partial return to a previous state. This is not merely a result of trivial features of spiking activity, such as simple temporal covariance in spike rates, or of rate changes that simply return to a previous level without a rotational component (see: “Controls confirming organized rotational dynamics”, below) .

To isolate the rotational dynamics, we used rotational principal components analysis (jPCA). jPCA identifies axes that capture the dominant rotational components of input data (Churchland et al., 2012; Gao et al., 2016; Jiang et al., 2020). We examined the rotational component in the spiking activity for every working memory condition (the two sample objects and two locations they appeared). To quantify the strength of rotational dynamics, we examined the proportion of variance in the data’s *dynamics* (first temporal derivative of spike rates) explained by rotational components. This metric, which we refer to as R^2^, is what jPCA axes are ordered by (Churchland et al., 2012), analogous to the role of variance explained in the *raw* data in traditional PCA. Because we wanted to examine the saccade-distractor’s overall effect on prefrontal activity, we performed jPCA on spiking data without removing the average activity across working memory conditions. We found that jPCA captured higher explained variance in the dynamics of the full spiking data (R^2^ = 0.55 ± 0.01)—indicating a much stronger rotational component—than in spiking data with the average activity removed (R^2^ = 0.24 ± 0.005, P < 0.0001; Supplementary Fig. 1a,b). Thus, all further analyses are performed on the full spiking data, without the mean activity removed.

The saccade-distractor induced rotational dynamics in population spiking on correctly-performed trials (Fig. 2a). Similar effects were observed in both the “sending” and “receiving” hemispheres (Fig. 2a, left vs right), in both NHPs (Fig. 2a, insets), and in both the dlPFC and vlPFC (Supplementary Fig. 1c). Thus, for further analysis, we initially analyzed data separately for hemispheres, areas, and NHPs, and then pooled results across them. All reported summary results reflect these pooled results, unless otherwise noted.

**Figure 2.**
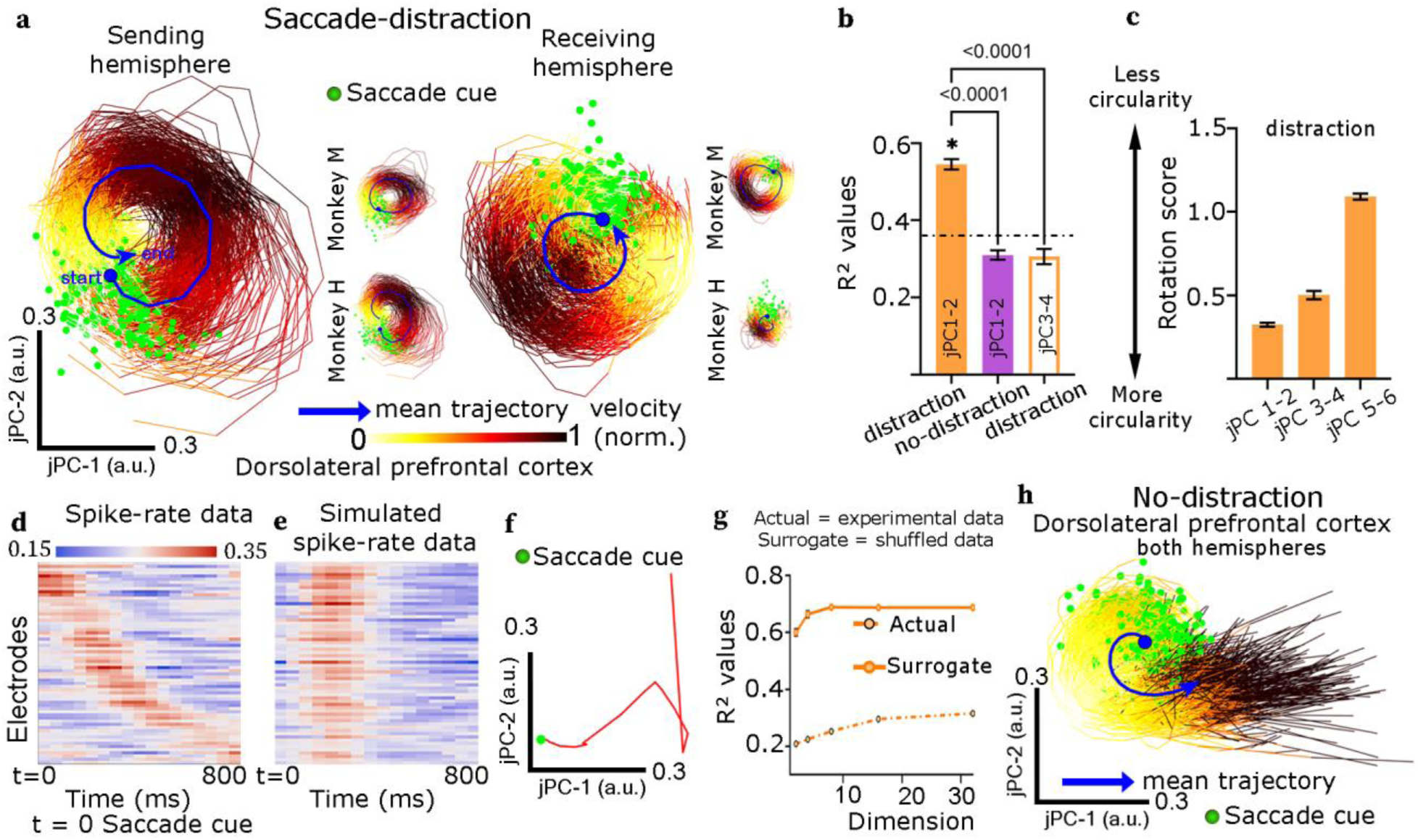
Saccades during working memory induce rotational dynamics (a) Projections of neural data on the top two jPCA components (jPCs) following the saccade (Delay 2) for both prefrontal hemispheres (sending = contralateral to the sample, receiving = ipsilateral to the sample) for the dlPFC (vlPFC results shown in Supplementary Fig. 1). Each of the large plots contains 220 trajectories, corresponding to the four working memory conditions ⨉ 55 sessions across two NHPs. Projections corresponding to individual NHPs are shown in the smaller inset plots. Each trajectory was color-coded based on the normalized linear velocity (normalized arc length traversed at each timestep). For clarity, trajectories were aligned within each plot. Differences in trajectory starting points between plots are therefore arbitrary (see Methods). Individual trajectories start from the mid-delay saccade cue (green circles) and span the entire Delay 2 (800 ms). The dark blue line indicates the mean trajectory (blue circle: saccade cue). (b) Proportion of variance explained in the data dynamics (R^2^) of fits of the top two jPCs, and the 3rd and 4th jPCs, pooled across both hemispheres and both PFC areas (dlPFC and vlPFC). R^2^ values were higher, indicating a stronger rotational component, for the top two jPCs than the third and fourth components. There was also a better fit (stronger rotational component) on trials with than without a saccade-distraction (P < 0.0001; paired t-test) for the top two jPCs. The dashed line indicates the criterion R^2^ value for significant rotational dynamics (*p* < 0.05, shuffle test; see Methods). (c) Rotation scores (reflecting trajectory circularity) of all trajectories combined for both hemispheres for the top three jPC planes during Delay 2. A score of zero indicates a perfectly circular arc. (d) Spike rate data sorted by latency to peak spike rate. (e-f) Simulated data with no sequential pattern produced non-circular trajectories in the subspace of top two jPCs. (g) R^2^ estimates of fits of the jPCs to the actual data and shuffled surrogate data. The surrogate data was generated by the TME algorithm, where all the dimensions—time, neuron and condition—were shuffled, but lower-order statistics along each of these dimensions were preserved. R^2^ was much higher in the actual data, indicating rotational dynamics could not be explained by simple features of the data. (h) Projections onto the top two jPCs for no-distraction trials.

We tested if there were subpopulations of neurons with distinct temporal activity profiles, which might make differential contributions to the rotational dynamics. We examined the distribution of peak activation times across all single neurons in our data. This revealed a smooth progression of peaks across time, with no evidence of grouping into distinct activity profiles (see Supplementary Fig. 5). This suggests that there was no such clustering of neurons into discrete subpopulations, and that all neurons contributed in a qualitatively similar fashion to the rotational dynamics. Thus, for further analyses, we analyze the entire neural population as a whole.

To ensure rotational dynamics reflected a substantial component of the *overall* data itself—not just its dynamics—we also examined how much variance they explained in the raw data (see Methods). Note that, in contrast to the dynamics explained variance introduced above, there is no expectation that jPCA components will also be ordered by the variance they explain in the raw data. Substantial raw variance in the data was explained by the first four jPCA components (jPCs; 0.66 ± 0.002). The first two jPCs explained about a third of the overall variance in the raw spiking activity (0.29 ±0.02). The rotational dynamics were also most circular for the top two jPCs and became progressively less circular for higher-order jPCs (Fig. 2c, see also Supplementary Fig. 1c,d). Thus, to isolate the strongest rotational dynamics in the data, we focused on the top two jPCs for further analyses (Churchland et al., 2012).

Rotational dynamics could be due to sequential activation of neurons in response to the distractor (Kuzmina et al., 2024; Liu et al., 2025). In fact, we observed this. Following the distractor, prefrontal neurons showed a sequential pattern with continuously offset peak latencies (Fig. 2d; see also Supplementary Fig. 2). jPCA effectively captured this temporal sequential structure. We reconstructed the spike rate data using only the top two jPCs and plotted it with the same neuronal ordering as in the actual data (Supplementary Fig. 2a). The reconstruction from the top two jPCs showed a similar sequential structure as the actual data (Fig. 2a). This was not the case for higher-order jPCs. When we performed the complementary analysis, reconstructing the data with the top two jPCs removed, the sequential structure was eliminated (Fig. 2a). Thus, we confirmed that state-space rotations corresponded to an orderly temporal progression of spiking activity. But this sequential activation was not arbitrary. In fact, our analysis showed that this orderly progression returned the spiking back to a state similar to that before distraction. We address this in the section “Properties of rotational dynamics” (see also Supplementary Fig. 3, 4). Note also that this progression only demonstrates a temporal sequence, and does not necessarily imply any spatial ordering. We will return to this issue later.

### Controls confirming organized rotational dynamics

We conducted several controls to confirm that the state-space rotations were due to organized population-level dynamics and were not due to simpler features of the data.

1. Rotations are not consistent with a simpler model of distractor-induced dynamics

An alternative model is that the distractor induces a phasic offset of population activity in state-space that simply returns along the same trajectory, perhaps due to simple temporal decay.This would correspond to the distractor inducing a brief burst of spikes that is simultaneous across neurons, rather than the ordered sequence we observed. To simulate this “out-and-back” situation, we shifted the activity time course of each neuron so that all of their peaks aligned in time (Fig. 2e). This eliminated the sequential pattern while otherwise preserving the statistics of individual neurons in the real data. When jPCA was performed on this temporally-aligned out-and-back data, there was no evidence of rotational state-space dynamics (R^2^ = 0.23 ± 0.007, Fig. 2f). This indicates that our results could not be explained by a simpler alternative model of distractor-induced dynamics that corresponds to non-sequential spiking activity.

1. Rotations reflected population-level patterns rather than simple features of spiking

To confirm that rotational dynamics are not simply a result of any low-level features of the data, we generated randomly shuffled datasets designed to match the actual spiking data’s mean and covariance across time, neurons, and working memory conditions. We used two different shuffling algorithms, corrected Fisher randomization (CFR) and tensor maximum entropy (TME) (Elsayed & Cunningham, 2017). This destroyed all higher-order joint statistics reflecting spatiotemporal population spiking patterns, including the sequential activation patterns described above, but preserved the mean and covariance of the spiking data over time, neurons, and task conditions. We used R^2^ values to measure the proportion of variance in the data dynamics explained by the jPCs (Fig. 2g; see Methods).

The shuffled datasets did not show rotational dynamics. The R^2^ of the jPCA fits was significantly less than that of the actual data, regardless of the number of jPCA dimensions considered (shown for TME in Fig. 2g, see Supplementary Fig. 6a,b for CFR and other variants of TME). This indicates that rotations reflect a population-level pattern not predicted from simpler features like cross-temporal or cross-neuronal correlations in the spike rates of individual neurons.

We used this same shuffling procedure to quantify whether population trajectories under various conditions contained rotational dynamics stronger than expected by chance (see “Significance testing of rotational dynamics” in Methods). We repeated the random shuffling to estimate a null distribution of R^2^ values expected by chance, when there is no true rotational structure in the data. The 95% percentile value of this distribution (R^2^ = 0.36) was used as a criterion value for significant population rotational dynamics in the saccade-distractor task.

1. Rotations were stronger after the saccade compared to no-saccade control trials

We examined the same time interval (corresponding to Delay 2) on trials in which there was no mid-delay saccade. R^2^ values reflecting the variance in the data dynamics explained by the top two jPCs were much higher (P < 0.0001; paired t-test) after the saccade (R^2^ = 0.55 ± 0.01; Fig. 2a,b) compared to analogous time periods in no-distraction control trials, which were not significantly different from chance (Fig. 2h,b; R^2^ = 0.31 ± 0.01). This indicates that there was no evidence for rotational dynamics when there was no saccade-induced perturbation of spiking activity.

1. Rotations were observed using methods not optimized to capture rotations

jPCA is designed to capture rotational dynamics. For a further control test, we looked for evidence for state-space rotations using measures not optimized for capturing rotations. As previously noted, we observed rotations using standard (non-rotational) PCA (Fig. 1b; Supplementary Fig. 7a). We also used t-SNE, a dimensionality reduction method that does not have any bias toward finding rotational dynamics (Arora et al., 2018). The top two t-SNE components did indeed reveal visual evidence of rotations in the data (Supplementary Fig. 7b). These results confirmed that rotational dynamics were present in the data, independent of using jPCA. However, we used jPCA for most of our analyses because it isolates rotational dynamics better than other methods

1. Rotations were not caused by “phantom oscillations”.

Temporal smoothing of data can induce oscillatory patterns in PCA (or jPCA) components even with no actual oscillatory dynamics in the data, a phenomenon called “phantom oscillations” (Shinn, 2023). As a control, we generated surrogate datasets that had the same PCs and associated data variances as the actual data, but were otherwise random (see “Surrogate data generation with oscillatory PCs” in Methods). This surrogate data showed no rotational trajectories (Supplementary Fig. 7c). Further, we examined the covariance matrix of data during Delay 2 (Supplementary Fig. 4). Covariance matrices reflecting “phantom oscillations” are expected to show only a single thick stripe along the main diagonal (Shinn, 2023). In contrast, in our data covariance matrix, we observed additional off-diagonal structure reflecting the return of activity to a similar state to before the distractor. These controls indicate that phantom oscillations induced by jPCA cannot explain our results.

1. Saccades in the absence of working memory requirements did not induce rotations

We wanted to rule out the possibility that rotational dynamics are induced by any saccade, whether or not there were working memories that needed protection against distraction. Thus, we examined situations where the NHPs made saccades with no working memory demands. We examined the initial fixation period (see Methods, and Supplementary Fig. 8), following the saccade to the fixation dot that initiated the trial. No working memory was involved during this period because the sample had yet to be seen. We also examined the behavioral response period, following the saccade to indicate the NHPs’ behavioral choice, when the working memory no longer needed to be retained.

We observed no significant population-level rotational dynamics during either of these periods (post-fixation: R^2^ = 0.15 ± 0.01, P = 1, one-sided t-test; post-choice: R^2^ = 0.35 ± 0.016, P = 0.53, one-sided t-test, Supplementary Fig. 8). This suggests that saccades alone are not sufficient to induce rotational dynamics. Rotations were only observed when working memories were retained across a saccade, and thus recovery from the saccade-induced perturbation was required.

### Properties of rotational dynamics

Following the saccade, neural trajectories completed approximately a full 360° rotation by the end of the memory delay (Delay 2, total rotation = 364.5 ± 3.87°). The angular rotational velocity peaked around the time of the saccade and then gradually decreased over the remainder of the delay (Fig. 3a). Similar effects, albeit weaker as expected, were found for the 3rd and 4th jPC components shown in Fig. 2b (open bar) and Supplementary Fig. 9a,b.

**Figure 3.**
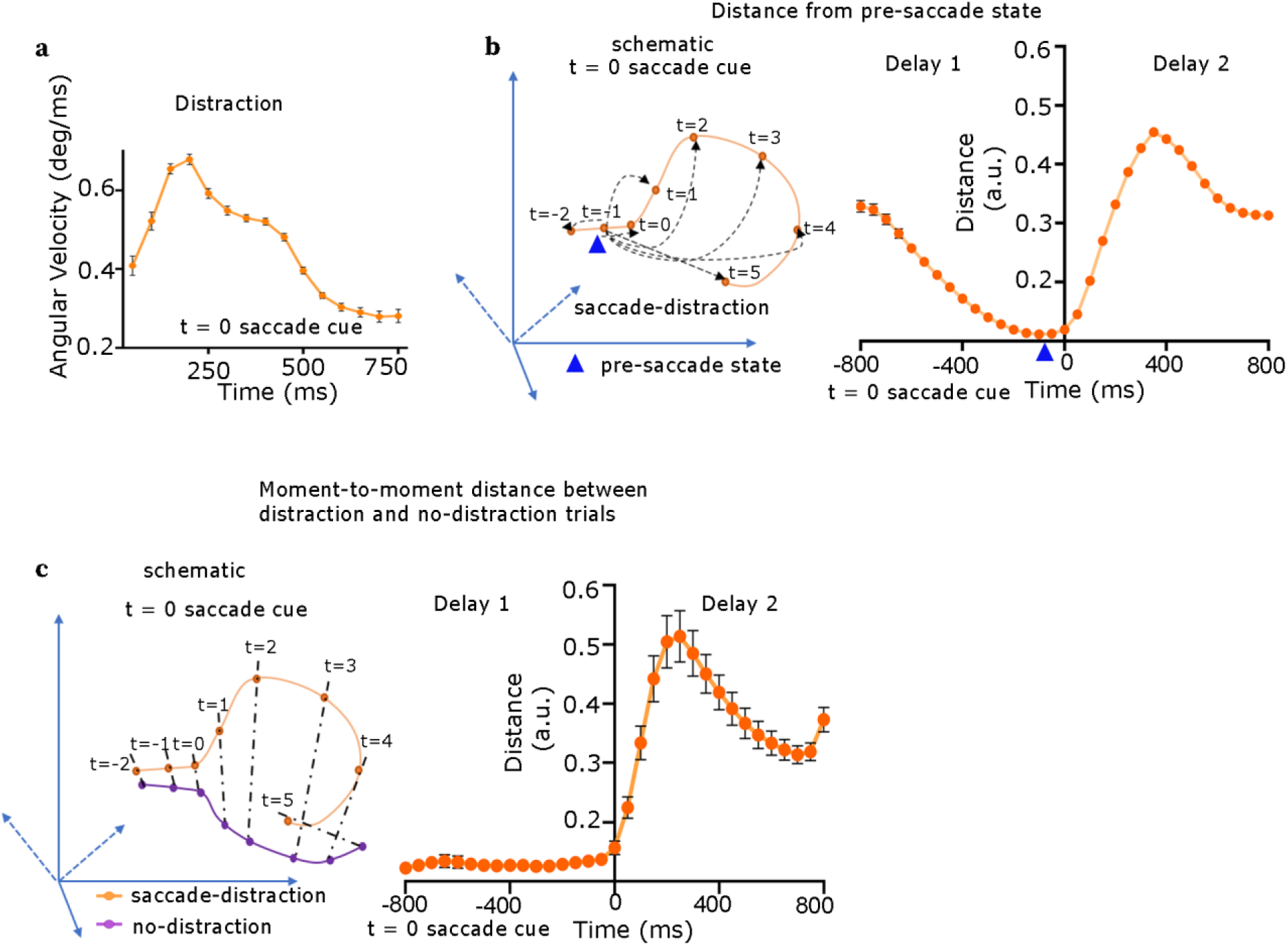
Properties of rotational dynamics (a) Angular velocity as a function of time for saccade distraction trials. (b) We computed the distance between the population spiking activity states at each post-saccade time point and the pre-saccadic baseline (blue triangle; see Methods for details). Cross-validated distances increased after the saccade-distractor, then decreased again, consistent with a partial recovery. (c) We computed distances between corresponding time points within neural trajectories on saccade and no-saccade trials. Distances initially increased after the saccade, then decreased, again consistent with a partial recovery.

The 360° rotation suggested that spiking patterns were returning to a state similar to that before the disruption. To determine the degree of similarity, we examined higher dimensions than 2D. It is possible that a 360° rotation seen in 2D (Fig. 2a) could be a helix in higher dimensions. If so, that would not return spiking to the exact same state even after a complete rotation. We found that this was the case. As a result, the 360° rotation returned spiking to a similar but not the exact same state.

To determine this, we plotted the distance between population spiking states before and after the saccade cue over time in a higher-dimensional space that no longer isolated the rotational component (Fig. 3b and see “Distance between a pair of states” in Methods). This distance of every state was measured relative to the state just prior to the saccade cue (marked with a blue triangle in Fig. 3b). Naively, distances might increase away from the initial state simply due to the effects of uncorrelated noise. To account for this, and make all values statistically comparable, we computed distances cross-validated across trials (see Methods). The distance, by definition, was lowest just before the saccade cue at time = 0 (the “pre-saccade state”). After the cue, the distance between the current state and the pre-saccade state at first increased, peaking at the middle of Delay 2 (400 ms from the saccade cue). But then it decreased, moving back toward the pre-saccade state but not reaching it. Instead it reached a state that was almost halfway back to the pre-saccade state. Thus, the rotations did not fully return spiking to the same state prior to the saccade (percentage of recovery = 40.06 ± 0.02%).

We also computed the distance between population states at the same time points within trials with a saccade distractor and the control trials without a saccade. The distance between population trajectories in the distractor vs no-distractor trials were close to zero before the saccade cue appeared on distractor trials. Then, after the cue, their distance increased, reflecting sharp differences in spiking patterns. Then their distance returned to a level about halfway to their pre-saccade distance (Fig. 3c). These results indicate that the observed rotation dynamics reflect a partial recovery that returned population spiking close to, but not identical to, its pre-distractor state.

### Rotations were less complete before errors

The saccade disrupted task performance (causing a 9.77 ± 0.07% drop in correct choice, see above). On error trials, the trajectories failed to complete a full rotation (in 2D) before the end of the delay (average = 333.5 ± 2.85°), unlike correctly performed trials (364.5 ± 3.87°, Fig. 4a,b, P < 0.0001, one-sided paired t-test; see also Supplementary Fig. 1c). Thus, the trajectories did not converge to the pre-saccade state. In contrast, the population mean spike rate did not differ between correct and error trials (P = 0.348, one-sided paired t-test). On error trials, the angular velocities started slower (Fig. 4c), and their trajectories were less circular and more elliptical (Fig. 4d). These results show that errors coincided with atypical rotational dynamics. They are consistent with errors resulting from a failure of rotational dynamics to return neural activity to close to its pre-distractor state.

**Figure 4.**
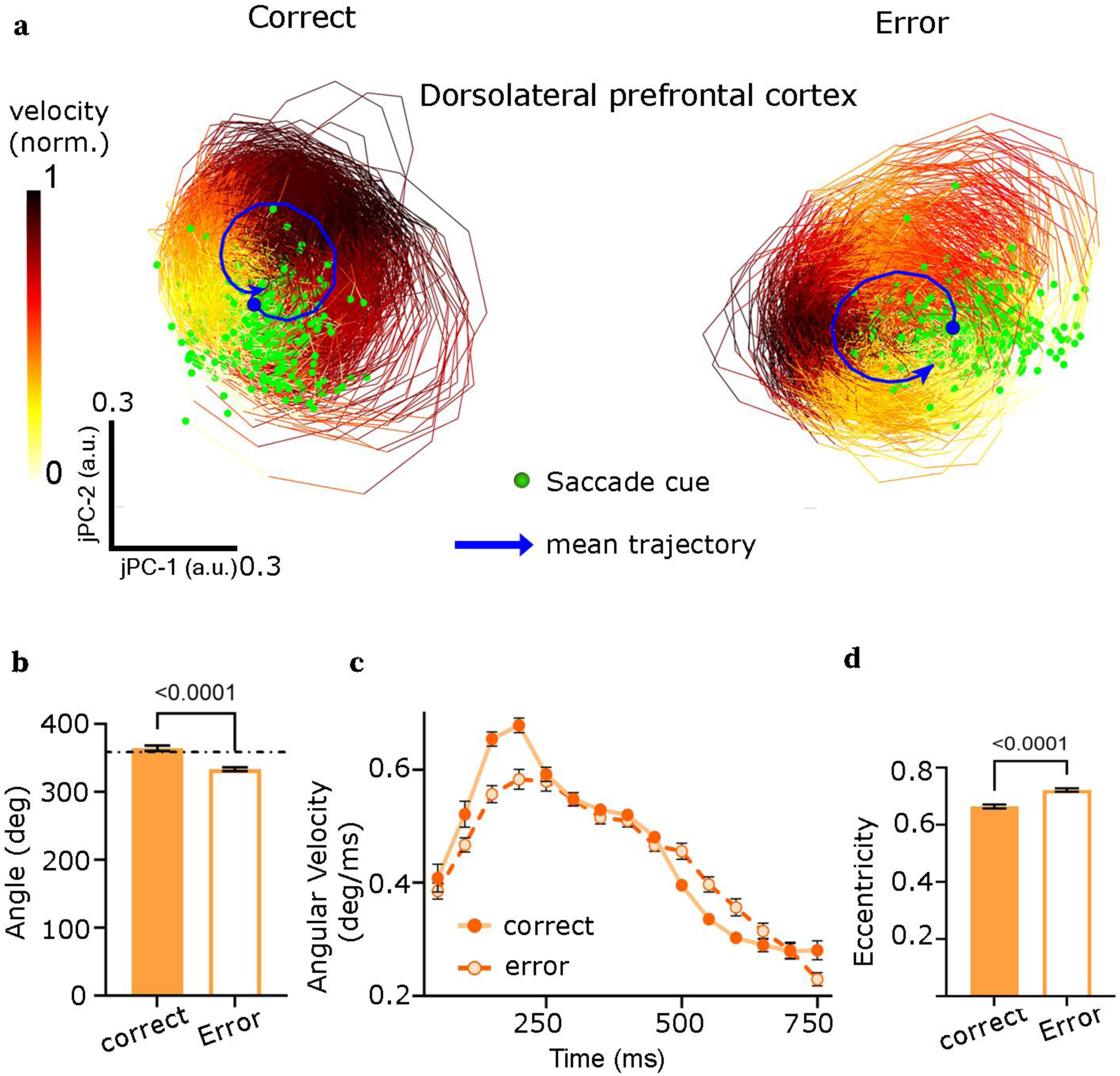
Behavioral effect of rotational dynamics. (a) Projections of dlPFC data onto top two jPCs during Delay 2 for all four working memory conditions for behaviorally correct (left) and error (right) trials. (b) Total angle of rotation for correct and error trials (differences in angles: P < 0.0001, t = 7.75, df = 54, paired t-test). The dashed line indicates 360 degree rotation. (c) Angular velocity during correct and error trials. During the 0-200 ms after saccade cue onset, angular velocity between correct and error trials was different (P = 0.0065, paired t-test). There was also a difference late in the delay (600-800 ms, P = 0.027, paired t-test). (d) Eccentricity of the correct and error trials. Error trials also had significantly more elliptical trajectories than correct trials.

As a further test, we examined whether neural coding of information held in working memory was stronger after fuller rotations. We fit linear classifiers to decode the position and identity of the object held in memory from population spiking activity during Delay 2, after the mid-delay saccade. We quantified the trial-by-trial strength of decoded information using the classifier posterior probability, averaged over the last half (final 400 ms) of Delay 2 (see “Angle of rotation and posterior scores” in Methods). We found that the angle of rotation was positively correlated with decoded information strength, for both objection location (*r* = 0.78, *p* = 0.0014, t-test) and object identity (*r* = 0.45, *p* = 0.012; t-test, Supplementary Fig. 10). These results suggest that fuller rotations result in stronger coding of information held in working memory.

### Rotations were also induced by visual stimulation

We next asked whether visual distraction would induce the same rotational dynamics as a saccade-distractor. We used another visual object working memory task (Fig. 5a). This time, a visual distractor (instead of a saccade) appeared in the middle of the memory delay. There were five different delay lengths (1, 1.41, 2, 2.83, and 4 s), randomly interleaved. The distractor always appeared in the middle of the delay. This means that the delay length before the distractor was unpredictable, but once the distractor appeared, the NHPs could predict the remaining time in the delay. We again began by focusing on the time that included the distractor and remainder of the memory delay (Fig. 5a, “Delay 2”).

**Figure 5.**
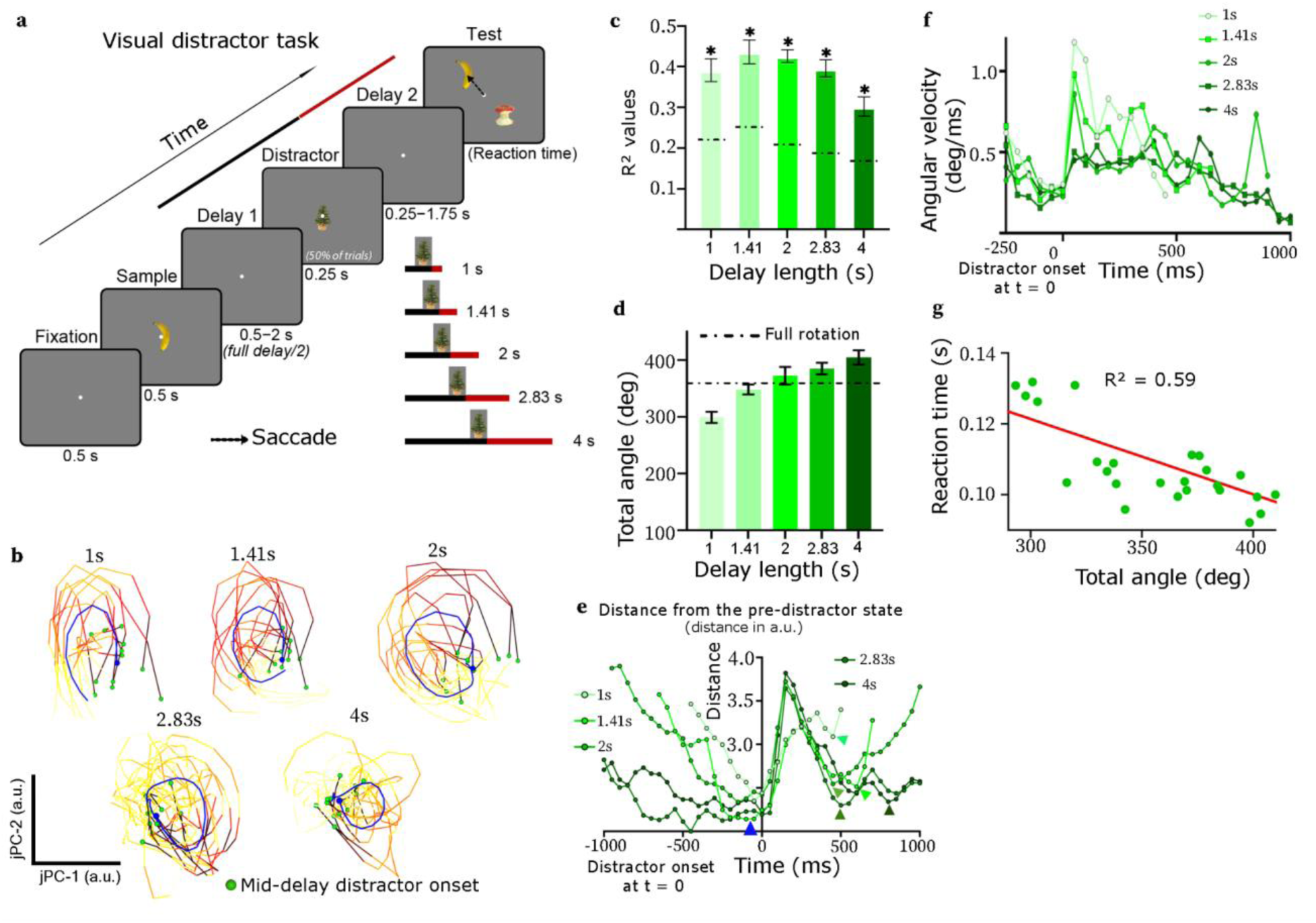
Rotational dynamics were also induced by visual distractors. (a) A delayed-match-to-sample working memory task with a visual distractor. Inset: The distractor always appeared in the middle of the variable-length delay, so the remaining post-distractor delay was predictable. (b) Projections onto the top two jPCs (combined for both the left and right hemispheres) for five delay lengths. The length of the trajectory increased with the length of the delay. Color of a trajectory denotes the normalized linear velocity (arc length/time). (c) Proportion of variance explained by the rotational dynamics (R^2^) of fits of the top two jPCs to the data. The dashed line indicates the criterion R^2^ value for significant rotational dynamics for individual delays (*p* < 0.05, shuffle test; see Methods). (d) Total angle of rotation for each delay length. Longer delays resulted in longer rotations. The dashed line indicates 360 degree rotation (a full rotation). (e) Cross-validated distances from the pre-distractor state (blue arrowhead) for each delay length. Colored arrowheads indicate the post-distractor states closest to the pre-distractor state for corresponding delays. The neural state generally showed more complete recovery for longer delays. (f) Angular velocity of rotations. Angular velocity peaked within 200 ms after the distractor onset. Peak angular velocity was inversely correlated with the delay length (*r* = −0.8 ± 0.11, P < 0.0001). (g) Reaction time (RT) as a function of total angle. RT was inversely correlated with the total angle.

The visual distractor also caused rotational dynamics. Spiking data for Individual delays showed strong rotational dynamics, captured by the projections onto the top two jPCs (Fig. 5b). At each delay length, rotational dynamics were stronger than expected by chance (Fig. 5c). Longer delays resulted in larger total rotations (Fig. 5d, correlation = 0.73 ± 0.12, P < 0.0001). As expected from the relationship between rotation angle and delay length, the degree of recovery (how much spiking returned to the pre-distractor state) was also dependent on the length of the delay. Longer delays led to a more complete recovery (Fig. 5e). In Fig. 5e, we measured the cross-validated distance between the neural state at each time point and the state just before the disruption. We found partial recovery for shorter delays (percentage of recovery = 32.63 ± 0.07% for 1s, 51.83 ± 12.77% for 1.41s; see Methods) and near-complete recovery for longer delays (82.33 ± 0.06% for 2s, 81.63 ± 0.06% for 2.83s, 94.93 ± 0.02% for 4s; Fig. 5e, see Methods). These results suggest that neural trajectories require time to return (near to) their pre-disruption state.

In fact, rotational velocity depended on the animal’s expectation of delay length, as if it were speeding up to compensate for the expected shorter delays. Across all delay lengths, the rotational velocity was fastest immediately after the visual distractor, then slowed down over the remainder of the delay (Fig. 5f), similar to that after the saccade (Fig. 3a). However, just after the visual distractor (∼0-200 ms), the rotational velocity was fastest for the shortest delay (1 sec) and decreased in a graded fashion with increasing delay length (correlation coefficient = −0.8 ± 0.11, P < 0.0001). This was not due to using a longer temporal duration of analysis at longer delays. The effect persisted even when jPCA was performed within the same time window (0– 500 ms) across all delay lengths (correlation coefficient = −0.77 ± 0.16, P < 0.0001). This suggests an adjustment of the velocity of rotational dynamics to match the animal’s expectation of delay length, which was predictable after the distractor.

The angle traversed by visual-distractor-induced rotations also correlated with behavior, as we previously observed for saccade-induced rotations. In this task, the NHP made few choice errors—behavioral accuracy was 99.8 ± 0.002% across sessions. Thus, to relate rotational dynamics to behavior, we examined how they correlated with the reaction time to make an initial saccade to one of the test stimuli. In no-distractor trials, reaction times were nearly constant across delay lengths (*slope* = −0.0008, *p* = 0.247), suggesting there was no substantial memory decay over these relatively short delays (Supplementary Fig. 11). In contrast, on distractor trials, reaction times were slower overall (*p* = 0.0232, one-sided t-test), but speeded up with longer delays (*r* = −0.47 ± 0.02, *p* = 0.0189, t-test). This indicates the visual distractor did indeed impair behavior, and that the NHPs required some time to recover from distraction. There was also an inverse relationship between the total angle of rotation and the reaction time in distractor trials. Reaction times were faster when rotations completed a larger angle (correlation = −0.4 ± 0.017, P < 0.0001, Fig. 5g). To quantify the independent contributions of delay length and rotation angle to behavior, we regressed reaction times on both delay length and rotation angle across all delay lengths and sessions. We found that both delay length (p = 0.0007) and rotation angle (p = 0.005) were strong predictors of reaction time. Thus, independent of any effects of delay length, a larger total angle of rotation was associated with faster reaction times. These results suggest that successful recovery from distraction may depend on completion of a full cycle of population rotations.

### Traveling waves corresponded with rotations in population activity space

The organization of spiking into rotational dynamics raised the possibility that spiking might also be organized not just by time but also by space. In other words, the spiking patterns might have formed traveling waves across the cortical surface (Alamia & VanRullen, 2019; Bhattacharya, Brincat, et al., 2022; Bhattacharya, Donoghue, et al., 2022; Davis et al., 2020; Goldman et al., 1949; Huang et al., 2010; Lubenov & Siapas, 2009; Mohan et al., 2024; Moldakarimov et al., 2018; Muller et al., 2016, 2016, 2018, 2018; Patel et al., 2012; Riehle et al., 2013; Rubino et al., 2006; Takahashi et al., 2011, 2015; Y. Xu et al., 2023; Zanos et al., 2015; Zhang et al., 2018) . We observed a temporal sequence of spiking activity when neurons were sorted by their latency (Fig. 2d). But note that this result does not imply any spatial organization. It could correspond to a spatially random “salt-and-pepper” distribution in which there was no correspondence between a neuron’s ordinal position within the temporal sequence and its location within cortex. Here, we instead show that it had an orderly structure in space as well, which was consistent with a traveling wave of spiking activity moving across the cortical surface.

To examine this, we returned to the saccade-distractor task (similar effects were seen after the visual distractor as well, see Supplementary Fig. 12). To quantify traveling wave structure, we computed the principal direction of the spatiotemporal flow of spiking activity across each electrode array using an optical flow method (see Methods). We then measured its angular deviation, the absolute change in the angle of the principal direction over time. Data lacking traveling waves would be expected to have random, uncorrelated changes in principal directions across successive time points, and thus, on average, a large angular deviation. In contrast, traveling waves would be expected to have a smooth, orderly change in principal direction over time (black arrows in Fig. 6a), and thus a small average angular deviation. We observed small changes in the angular deviations of the principal directions over time, consistent with traveling waves (Fig. 6b, blue line). As a further confirmation, we used surrogate data generated by the TME (like the one in Fig. 2g) algorithm as a control (Fig. 6b, gray line). The shuffling disrupted the traveling wave structure in spiking activity, as indicated by significantly increased angular deviations (post cue-to-saccade: actual data = 36.6 ± 1.95°, surrogate data = 56.2 ± 2.5°, P < 0.0001). We also confirmed the presence of traveling waves using a method that did not depend on computation of optical flow. It instead directly quantified correlations between activation latency time and spatial distance between pairs of neurons (see “Alternative method for traveling wave detection: Correlation of activation latency with distance” in the Methods section and Supplementary Fig. 13) (Patel et al., 2012). The correlation was higher in the actual data than the shuffled surrogate data (actual data: 0.45 ± 0.005, surrogate data: 0.16 ± 0.001, P = 0.016, one-sided paired t-test). These results indicate that there were traveling waves of spiking activity across the cortical surface during the distractor tasks.

**Figure 6.**
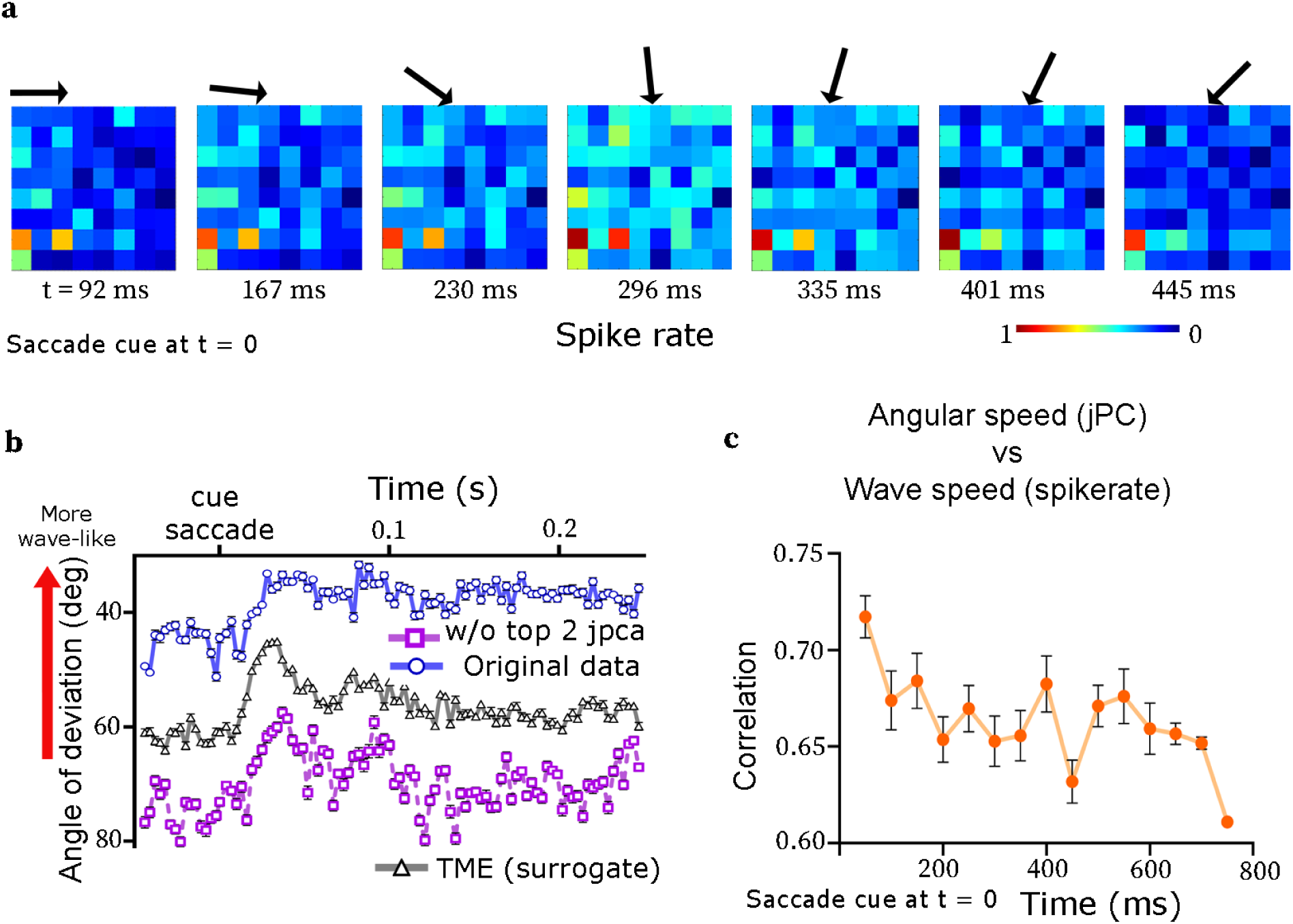
Rotational dynamics corresponded to traveling waves across the cortical surface (a) An example traveling wave in the trial-averaged spike rate. Arrows indicate principal direction of flow of spiking activity across the array. (b) Mean angular deviation of the principal direction of spike traveling waves across sessions (blue line). The same data is shown after subtracting the top two jPCs (magenta line) and after shuffling the data by TME algorithm (black line). (c) Moment to moment correlation between the angular speed of state-space rotations (from the projections in the subspace of top two jPCs) and traveling wave speed (principal magnitude; see Methods).

The traveling waves were not induced by the location of the fixation point sweeping across neural receptive fields during the saccade. The duration of the saccades–where such a sweeping effect would be possible–was only about 20 ms, whereas the wave lasted the entire duration of Delay 2 (800 ms). Given that prefrontal sensory responses typically decay after ∼150 ms after stimulus offset (e.g. Brincat et al. 2021 Fig. 2), they seem unlikely to induce such long duration traveling waves. Further, in the visual distractor task, the distractor image was shown at the fovea during fixation, and thus there was no possible “sweeping” effect. Nevertheless, we observed traveling waves that lasted from 500 ms to 2s based on the delay duration (Supplementary Fig. 12).

The presence of both traveling waves across cortex and population state-space rotations after the saccade distractor raised the possibility of their correspondence. To test this, first we measured moment-to-moment correlation between the speed of the spiking traveling wave and the angular velocity of the trajectory (see Fig. 3a) in the subspace of the top two jPCs (see Methods). We found a high correlation (e.g. 0.7 ± 0.03 at 100 ms after the saccade cue; Fig. 6c). The correlation dropped towards the end of Delay 2 when rotational dynamics also weakened, but remained well above chance (0.63 ± 0.01, 100 ms before the end of Delay 2). As expected from the correlations, traveling waves, like state-space rotational dynamics, were fastest after the saccade cue and then slowed over the memory delay (Fig. 6c).

To confirm this relationship, we demonstrated that removing the projections of the spiking data onto the top two jPCs also disrupted the traveling wave. We removed the first two jPC projections from the spiking data, then recomputed its principal directions. This resulted in a significant increase in angular deviation, indicating a loss of traveling wave structure in the spiking activity (Fig. 6b, magenta line, difference in angular deviation = 31.32 ± 0.45°, P < 0.0001, paired t-test). These results suggest the possibility that there is a correspondence between rotational dynamics in spiking activity and traveling waves of spiking across the surface of cortex.

## Discussion

Neural spiking in the PFC showed a rotational component in population activity state-space after distraction. This reflected an organized progression of spiking patterns that partially returned spiking to a state more similar to that before a distraction. Rotations were fuller when the task was correctly performed. We also found evidence that state-space rotations corresponded with traveling waves rotating across the surface of the cortex. This suggests a possible link between traveling waves and state-space neural dynamics. These dynamics could correspond to a crucial brain function: dynamic stability (Kozachkov et al., 2020). Despite variation and noise in neural activity, the brain has to converge to a stable, reproducible sequence of states and return to the proper “track” when perturbed, a function referred to as “dynamic stability”(Buonomano & Maass, 2009; Ju & Bassett, 2020; Kozachkov et al., 2020). Broadly similar results were seen with both saccade and visual distractors, suggesting they may reflect a common mechanism for cortical recovery from all types of perturbation.

Rotational dynamics seem to be a common motif in the cortex. They have been reported in the motor, premotor, somatosensory, visual, parietal, and prefrontal cortices (Aghagolzadeh & Truccolo, 2016; Aoi et al., 2020; Churchland et al., 2012; Diomedi et al., 2021; Gallego-Carracedo et al., 2022; Jiang et al., 2020; Kalidindi et al., 2021; Kraus et al., 2013; Lara et al., 2018; Lebedev et al., 2019; Mollazadeh et al., 2014; Suresh et al., 2020; Sussillo et al., 2015; Townsend et al., 2015; Townsend & Gong, 2018). Rotations can, in essence, return spiking to a state near to that before the perturbation that induced them, much as working memory representations are temporarily lost and then reinstated following distraction (Derrfuss et al., 2017; Mallett & Lewis-Peacock, 2019; Yoon et al., 2006) . The rotational dynamics could arise from a wide range of anatomical connectivity patterns(Churchland et al., 2012; Davis et al., 2021, 2024; Kalidindi et al., 2021; Suresh et al., 2020). Population-level state-space dynamics seems to be under top-down influence. They can change speed with task demands(Wang et al., 2018). We observed a similar property in state-space rotations. When the expected remaining memory delay was shorter, population activity rotated with higher velocity, while longer rotations had slower velocity.

Rotational dynamics do not necessarily imply a complete “reset” back to a previous state. Mathematically, the difference between the final state and the initial state can fall into the null space of the first two jPCA axes after the rotation. That is, there can be differences along dimensions orthogonal to the jPCA axes, as in a helix. Our results were consistent with this. Thus, they reflected only a partial recovery back toward a previous state. Partial recovery may also reflect the time available to recover from a perturbation. When the post-distractor delay was shorter, recovery was less complete. With longer post-distractor delays, population activity made a near-complete recovery to its pre-distractor state.

Traveling waves of activity across the surface of cortex are another emergent phenomenon(Bhattacharya, Brincat, et al., 2022; Bhattacharya, Donoghue, et al., 2022; Huang et al., 2010; Muller et al., 2016; Y. Xu et al., 2023). The state-space rotations we observed seemed to correspond to traveling waves of spiking activity. This is consistent with previous work linking rotational dynamics to sequential neural activation(Kuzmina et al., 2024). Like state-space rotations, traveling waves are common across cortex (Benucci et al., 2007; Bhattacharya, Brincat, et al., 2022; Bhattacharya, Donoghue, et al., 2022; Gabay et al., 2018; Goldman et al., 1949; Huang et al., 2010; Kuzmina et al., 2024; Roberts et al., 2019; Y. Xu et al., 2023; Ye et al., 2023). Traveling waves have properties useful for a variety of functions (Alamia & VanRullen, 2019; Davis et al., 2024; Lubenov & Siapas, 2009; Muller et al., 2016, 2018; Pang et al., 2020; Patel et al., 2012; Takahashi et al., 2011), including regulating attentional scanning and tracking time and network activity (Fries, 2023; Martinius & Hoovey, 1972; Scolari et al., 2015; Silver & Kastner, 2009). Visual targets are better detected when traveling waves in the cortex are better organized(Davis et al., 2020). The theory of spatial computing shows how such ordered spatiotemporal changes in activity could underlie the control of spiking activity(Lundqvist et al., 2023). This suggests a relationship between traveling waves and state-space dynamics(Huang et al., 2010; W. Xu et al., 2019).

## Methods

### Behavioral paradigm

We employed two visual working memory tasks. In both tasks, there was a distraction in the middle of the memory delay, either a cued shift in gaze or an irrelevant visual distractor.

The saccade-distraction task (monkeys M and H) is shown in Fig. 1a. The trial began with 700 ms of central fixation. Then, a sample object appeared for 700 ms to the right or left visual hemifield, either 3.4° above or below the center of the screen. The sample was one of two objects from a commercial photo library (Hemera Photo-Objects). The task required memory of the sample object and location over a memory delay period of 1600 ms. On 50% of the trials, a gaze shift was instructed by moving the fixation point across the vertical midline to the opposite side (“saccade”, Fig. 1a). This gaze shift moved the remembered retinotopic location of the sample, being held during the working memory delay, to the opposite visual hemifield. At the end of the delay, a test object appeared for 400 ms. The monkeys were required to saccade to it if it did not match the remembered sample in either object identity or upper/lower location. If the test was identical to the sample, they withheld response. Then, after a 100 ms blank period, a non-matching test object was always shown, which required a saccade. Response to the non-match was rewarded with juice, followed by a 3.2 s inter-trial interval. Note that the 800 ms delay between the mid-delay and response saccades is much longer than the typical ∼200–250 ms interval between successive saccades in natural vision (Henderson & Hollingworth, 1999). Thus, it seems unlikely that any interference between the two saccades in our task contributed to behavioral errors.

In another “visual-distractor” task, an NHP (monkey H) performed a delayed-match-to-sample task with a mid-delay visual distractor (Fig. 5a). The trial began with 500 ms of central fixation. A sample object appeared for 500 ms at the center of the screen, followed by a variable-length delay period. The duration of the delay was randomly selected from a logarithmically-spaced set ([1s, 1.41s, 2s, 2.83s, 4s]). In a random 50% of trials, a distractor object (from a different set of objects as the sample) was presented for 250 ms at the middle of the delay. During the test period, the NHP was allowed to freely saccade between test objects and then indicate the final choice by holding fixation on it for 1s.

All the stimuli were displayed on an LCD monitor. An infra-red eye-tracking system (Eyelink 1000 Plus, SR-Research Ontario, CA) continuously monitored eye position at 1 kHz.

### Electrophysiological data collection

For both tasks, subjects were chronically implanted in the lateral prefrontal cortex (PFC) with four 8×8 iridium-oxide “Utah” microelectrode arrays (1.0 mm length, 400 *μ*m spacing; Blackrock microsystems, Salt Lake City, UT), for a total of 256 electrodes. Arrays were implanted bilaterally, one array in each of the ventrolateral and dorsolateral PFC in each hemisphere.

Electrodes in each hemisphere were referenced to a separate subdural reference wire. Spiking activity was amplified, filtered (250–5000 Hz), and manually thresholded to extract spike waveforms. All threshold-crossing spikes on each electrode were pooled together and analyzed as multi-unit activity (MUA). We recorded 20 sessions from monkey M and 35 from monkey H for the saccade-distractor task, and 6 from monkey H for the visual-distractor task.

### Quantification and statistical analysis

We examined neural responses to a saccade within the left and the right hemispheres separately in saccade trials. The results were pooled based on whether the sample was initially in the hemifield contralateral (sending) or ipsilateral (receiving) to the recorded hemisphere. The analysis was carried out using baseline-normalized spike rate data during Delay 2 (800 ms) in saccade trials (Fig. 1a) and at an identical time period (800 ms) in no-saccade trials, as controls to the saccade trials. In the experiment with a mid-delay visual distractor, we performed analyses separately on each hemisphere and the results were pooled to perform statistical tests. For both experiments, individual sessions were treated as observations. All processing and computational analysis were performed using MATLAB R2021b (Mathworks Inc., Natick, MA). The plots were generated using Prism (Graphpad software, MA).

### Preprocessing

Spike rates were computed by binning the spike timestamps in non-overlapping 50 ms windows. For the estimation of wave characteristics we used overlapping sliding time windows of 50 ms, each separated by 3 ms. The spike rates were square-root transformed to render the Poisson-like distribution of the spike rates to an approximate normal distribution. Further, with large samples (in our case, n = 55 sessions) the underlying distribution approaches the normal distribution (central limit theorem). This allowed us to use parametric statistical tests.

### Mean activity analysis

To normalize out any differences in activity between the neurons in all the working memory conditions, the spike rates were z-scored to the fixation baseline. We took a 500 ms time window before the sample onset and computed spike rates. The spike rates were then mean-pooled separately for each neuron and for each condition for each array location. The mean and standard deviation across all within-condition trials were computed. For each neuron and condition, these statistics were then used for z-scoring spike rates across all trials. We used z-scored spike rates for further analysis.

### Rotational PCA (jPCA) to capture rotational dynamics

To identify the population-level rotational dynamics from spiking data, jPCA was used. jPCA is a dimensionality reduction algorithm that finds the strongest rotational components of the neural activity(Churchland et al., 2012). Trial-averaged z-scored spike rate data with dimensions (neuron × timebin) was used as input to jPCA. For each neuron *i*, we computed the range of its spike rate along the time dimension and performed soft-normalization as

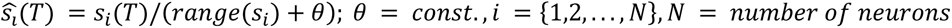

Soft-normalization rendered the neural responses to have a similar range of firing activity. We used 0.5 as the constant in all of our analysis. Most of our analysis focused separately on those aspects of neural responses that were common to, or that differed across, working memory conditions. We refer to these as condition-independent and condition-dependent respectively. In condition-dependent analysis, the across-condition mean was subtracted off the data from each condition.

At each level, the soft-normalized data were put together or stacked to compile an augmented data matrix, = [*X*_1_ *X*_2_ . . . *X*_*M*_]; *X*𝜖*R*^*N*^ ^×^ ^*MT*^. Here, M is the number of working memory conditions; N is the number of neurons; T is the number of time bins. X was then projected onto the subspace of its top six principal components (PCs), reducing X to a low dimensional data X_low_ of size (6 × MT). This step ensured that when jPCA was subsequently applied, it revealed only patterns of activity that reflected substantial data variance. Six principal components preserved at least 90% of the data variance.

To compute jPCA, an antisymmetric state-space matrix, D_skew_ 𝜖*R*^6^ ^×6^ was computed by fitting a linear model, 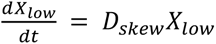. Quality of the fit was assessed by R^2^ values. R^2^ denotes the ability to linearly predict 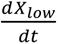 from X_low_, which, in turn, indicates the strength of rotational dynamics in neural responses. By definition, D_skew_ has imaginary eigenvalues, capturing rotational dynamics. The two eigenvectors U_1_ and U_2_ of D_skew_, corresponding to the two largest eigenvalues (one eigenvalue is the conjugate of the other) formed a plane that captured the strongest rotational structure in the data. U_1_ and U_2_ were complex-valued vectors. The jPCs were obtained as jPC_1_ = (U_1_ + U_2_)/2 and jPC_2_ = j(U_1_ − U_2_). The jPC projections were obtained as [jPC_1_;jPC_2_]*X_low_. The jPCs spanned the identical subspace to that of the six PCs, but with a basis optimized to identify rotational dynamics. We computed jPCA on each session separately. In each session, the jPCA algorithm yielded four projections corresponding to four working memory conditions ((obj1,UP), (obj1,DN), (obj2,UP), (obj2,DN)) separately for the saccade-distractor data and the no-distractor data.

To optimally align condition-independent trajectories across sessions for visualization, we first computed an average trajectory of all the working memory conditions. We took the average trajectory at session 1 as a reference. We obtained the best rotation, say 𝛩*i*, to fit between the average trajectory of the *i*^*th*^session and the reference. The trajectories of individual conditions were then rotated by 𝛩*i* to align with the reference. Finally, an average trajectory across all sessions was computed as the mean of the aligned trajectories. We performed these steps separately for the saccade-distractor data and the no-distractor data. This procedure was used only for the purpose of visualization (e.g. Fig. 2a). No computational analysis was performed on the rotated trajectories. Note also that this alignment was computed separately for each plot, hence their different—but arbitrary—starting points and global orientations.

#### jPCA in variable-delay visual-distractor task

In the experiment with a mid-delay visual distractor, it was necessary to compute jPCA in multiple phases due to the variable length of the post-distractor delay period. The shortest delay duration was 1 s. To calculate the jPC projections for 500 ms from the distractor onset, we considered all five delay durations (1, 1.41, 2, 2.83, and 4 s) as conditions and analyzed the trajectories for the 1 s delay only. To compute projections for larger delays, we progressively dropped one delay-length condition from the analysis. For example, to analyze dynamics for 700 ms after distractor onset, we only included delay-length conditions ≥ 1.41 s. Note that, within each delay-length condition, the sample was taken from a set of eight images. We pooled trials across all the samples and trial-averaged separately for each delay length.

### Rotation feature computation

We computed the following features of rotational trajectories in our analysis; (1) angular velocity, (2) eccentricity, (3) rotation score, and (4) R^2^ values from D_skew_. These features were computed separately for individual sessions. We then computed statistics on these features across sessions (i.e. using sessions as observations).

#### Angular velocity

Angular velocity is the angle subtended by an arc over time relative to its center. Computing this first required estimation of the center of a trajectory arc. Because of varying arc lengths in consecutive timesteps, a rotational trajectory was irregularly sampled in time. Thus, prior to estimating the center, each trajectory was interpolated to have *Q* regularly-spaced points along its length. Each point has 2D coordinates in the plane formed by jPC_1_ and jPC_2_. The center of the trajectory was computed as 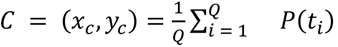 𝑃(𝑡); Here, t_i_ 𝜖[0,1] is the new parameter of the vector-valued trajectory *P*. We set *Q* = 50. Next, the angular velocity at each point of the original, non-interpolated trajectory was computed relative to *C*.

#### Eccentricity

Eccentricity quantifies the degree to which a trajectory is circular (low eccentricity) vs elliptical (high eccentricity). To compute the eccentricity of a trajectory, we fit an ellipse using a Matlab function (https://www.mathworks.com/matlabcentral/fileexchange/3215-fit_ellipse). Using the estimated major (*a*) and minor (*b*) axes, the eccentricity was computed as 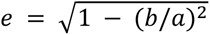. It is a real number in the range [0, 1], where zero indicates a perfect circle. It can be interpreted as a measure of the degree to which overall firing rates over time are relatively constant (eccentricity ∼ 0) or vary widely (eccentricity ≫ 0).

#### Rotation score

The rotation score also quantifies the degree to which a trajectory is circular, but against an alternative of any non-circular trajectory. It is based on the fact that, for a circular trajectory, the direction of the trajectory’s change over time is always orthogonal to the trajectory itself.

Rotation scores were computed using the relationship 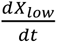. Let us assume that ^*X*^_*low*_ and 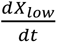 matrices contained *T* columns corresponding to *T* time points. We computed the angle, 𝜙(*i*) between 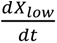(*i*) (trajectory change over time) and *X*_*low*_ (*i*) (the trajectory itself) at each timestep (*T* = *i*) on a trajectory. Each 𝜙(*i*) lie in [0, 2 ∗ 𝑝*i*). We computed the final rotation score at each time as the normalized deviation of 𝜙(*i*) from pi/2 radians: 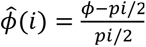. A rotation score of 0 corresponds to a perfect circle; a score of 1 corresponds to a trajectory making an instantaneous 180° radially inward hairpin turn; a score of −1 corresponds to a radially outward line. Trajectory rotation scores were summarized by their across-time means to obtain the final rotation score of the trajectory.

### Raw data variance vs dynamics explained variance

To compute the raw data variance captured by jPCs, at first, the actual spike rate data, *X*, was projected onto the top six PCs (see “Rotational PCA (jPCA) to capture rotational dynamics” section) to obtain a lower-dimensional projection of the data, *X*_*low*_. *X*_*low*_ was already mean-subtracted because we performed PCA. To obtain the jPC planes, we fitted the model 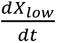<u>^=^</u>𝐷_𝑠𝑘𝑒𝑤_*X*_*low*_. Let the projection of *X*_*low*_ onto a jPC plane be *X*^_*low*_. The proportion of raw data variance of *X*^_*low*_ captured by the jPC plane was computed as 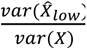.

In order to compute the rotational dynamics explained variance (R^2^), we used 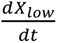. Let 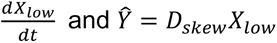 and 𝑌^ = 𝐷_𝑠𝑘𝑒𝑤_^*X*^_*low*_ be the estimate of 𝑌 from the model fit. Thus, the fitting error is 𝜀 = 𝑌 − 𝑌^ . Next, 𝑌 and 𝜀 were projected onto a jPC plane, resulting in 𝑌_*proj*_ and 𝜀_*proj*_, respectively. The dynamics explained variance of the data by the plane was computed as *R*^2^ = 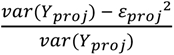. This is the same statistic reported in Churchland, et al. 2012.

#### Surrogate data generation with oscillatory PCs

To rule out that our results might be caused by “phantom oscillations” due to performing PCA on temporally smoothed data (Shinn, 2023), we generated surrogate data that had the same PCs and associated data variances as the actual data, including any PCA-induced “phantom oscillations”, but were otherwise random (Elsayed et al., 2016). Let *X* = [*X*_1_ *X*_2_ . . . *X*_*M*_] 𝜖 *R*^*N*×*MT*^be the trial-averaged spike rate data from an individual session, where *N* = number of neurons, *T* = total time of individual trial, *M* = number of working memory conditions. At first, *X* was soft-normalized(Churchland et al., 2012) and then only the duration during Delay 2 was considered from each *Xi*, forming the data 𝑌𝜖 *R*^*N*×*MT*𝑑^. Here *T*_𝑑_ is the time during Delay 2. Next, *Y* was mean-subtracted across time. We performed singular value decomposition (SVD) of *Y* to have *U* (left singular vectors), *S* (singular values) and *V* (right singular vectors). By formulation, *U* is also the set of eigenvectors of the covariance of *Y*. Each eigenvector contained evidence of oscillations.

To generate the surrogate dataset, we computed a matrix 𝑃𝜖 *R*^*N*×*MT*𝑑^, where each entry of *P* was randomly sampled from a uniform distribution with range [0,1]. *P* was mean-subtracted and then we computed the right singular vectors 𝑉_𝑃_ of *P*. We computed the surrogate data as *X*_𝑠_ = 𝑈 ∗ 𝑆 ∗ 𝑉^*T*^. *X*_𝑠_ has the same set of eigenvectors (principal components) and eigenvalues (component magnitudes) as of the actual spike rate data *X*.

### Surrogate data generation with preserved simple features

To generate surrogate data with the same marginal statistics as the actual data, we used the corrected Fisher randomization (CFR) and tensor maximum entropy (TME) methods (Elsayed & Cunningham, 2017). First, the data were randomly shuffled 100 times across either time (T), both time and neuron, (TN) or across all of time, neuron, and condition (TNC). This step does not preserve the primary feature statistics of the data (first-order and second-order marginal moments, i.e. mean and covariance, along the time, neuron, and condition dimensions). The CFR and TME methods do preserve such statistics, disrupting only higher-order statistics in the multivariate population data. Preservation of features can be accordingly done along the T or TN or TNC dimensions. We tested our results against surrogate data preserving statistics along time (T), time and neurons (TN), or across all three dimensions (TNC).

### Significance testing of rotational dynamics

To generate a null distribution reflecting the strength of rotational dynamics expected by chance, we computed R^2^ values by applying jPCA on the surrogate data, generated as described above. For each session, hemisphere, area, and surrogate generation method (CFR and TME), we generated random 100 surrogate datasets. Thus, each of these reflects the distribution of R^2^ values expected purely by chance, in the absence of any true rotational dynamics or other higher-order population-level structure in the data. We computed the 95th percentile of each of these distributions. We then averaged those values across sessions, areas, hemispheres, and surrogate generation methods. We used the average value (R^2^ = 0.36) as a criterion value to test for the presence of rotations. Only observed R^2^ values greater than the threshold (R^2^ = 0.36) were considered to have significant rotational dynamics (*p* < 0.05). In the Results, we refer to this as a “shuffle test”.

#### Distance between a pair of states

We computed the difference between pairs of neural states using the Mahalanobis distance between population patterns of spike rates. Because distances are non-negative, coupled with the considerable trial-to-trial variability in spike rates, distances would naively be positively biased. Even identical states could have non-zero distance due to being distinct samples of noisy data. To overcome this issue, we used a cross-validated Mahalanobis distance(Walther et al., 2016), where distances were computed between distinct subsets of trials.

In each session and for each condition, we randomly split the trials into two disjoint train and test groups with equal numbers of trials. This following procedure was run five times with random trial splits and then results were averaged. Let *X*_𝑡𝑟𝑎*i*𝑛_ = [*X*_1_ *X*_2_ . . . *X*_*M*_] 𝜖 *R*^*N*×*MT*^ be the trial-averaged spike rate data for the training dataset, where *N* = number of neurons, *T* = total time in each individual trial, *M* = number of working memory conditions. Similarly, let *X*_𝑡𝑒𝑠𝑡_ be for the test dataset. At first, *X*_𝑡𝑟𝑎*i*𝑛_ was soft-normalized and then only the duration during Delay 2 was considered from each *X*_𝑘_; 𝑘 𝜖{1,2, . . ., *M*}, forming the data 𝑌_𝑡𝑟𝑎*i*𝑛_𝜖 *R*^*N*×*MT*𝑑^. Here *T*_𝑑_ is the time during Delay 2. The same procedure was followed for the test dataset to obtain 𝑌_𝑡𝑒𝑠𝑡_. Next, we performed PCA and projected 𝑌_𝑡𝑟𝑎*i*𝑛_ onto the top ℎ principal components that collectively captured at least 95% of the data variance, which was ℎ = 10 for this data. Let this projection be 𝑍_𝑡𝑟𝑎*i*𝑛_𝜖 *R*^ℎ×*MT*^. Let the corresponding diagonal covariance matrix be 𝛴_𝑡𝑟𝑎*i*𝑛_𝜖 *R*^ℎ×ℎ^. The test data, 𝑌_𝑡𝑒𝑠𝑡_ were also projected onto those ℎ principal components as 𝑍_𝑡𝑒𝑠𝑡_𝜖 *R*^ℎ×*MT*^. We designated the average of the two time points immediately prior to the saccade cue as the pre-saccade “reference” state (Fig. 3b, blue arrow). We measured the distance of all other states (time points) to this reference. The “reference” state was taken from the test subset, and each compared time point was taken from the training subset . For the 𝑘^*th*^ condition 𝑘 𝜖{1,2, . . ., *M*}, let the pre-saccade “reference” state be 𝑠_𝑘_ ∈ *R*^ℎ×1^, and the compared state at time 𝑡 be 𝑧_𝑘_(𝑡) ∈ *R*^ℎ×1^. Then, the distance between each state and the pre-saccade reference was computed as 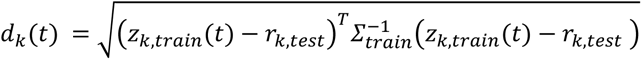. This was performed separately for each condition, hemisphere, and session, then the final result was averaged across them.

For the visual distractor experiment (Fig. 5a), we followed the same approach. We set the average of the two time points immediately prior to the visual distractor onset as the pre-distractor “reference” state (Fig. 5e). We computed the cross-validated Mahalanobis distance between each neural state and the reference state using the aforementioned method (Fig. 3b). We also separately computed the distance between corresponding timepoints within trials containing a distractor vs trials without a distractor (Fig. 3c). We quantified the degree of recovery in terms of the percentage of distractor-induced shift that was subsequently recovered. Let the distance value at the reference state be 𝑑_𝑟𝑒𝑓_, the peak distance from the reference state following the distractor be 𝑑_𝑚𝑎𝑥_, and the minimum distance from reference during the recovery by 𝑑_𝑚*i*𝑛_. We defined the percentage of recovery as 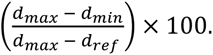.

### Sequential activation simulation

In order to understand how different population activity patterns relate to measured rotational dynamics, we simulated sequential neural activation patterns using time-shifted gaussian temporal activity profiles. The simulated data had the same dimensions as the actual spike rate data (neurons X time). For the saccade-distractor task, this was 64 neurons ⨉ 16 time points (separated by 50 ms). Each neuron’s activity across time was simulated as a gaussian pulse with standard deviation randomly selected from a uniform distribution between 50 and 200 ms. The gaussian mean (reflecting peak activity) for each neuron was set so they shifted in time continuously (discretized to the time sampling) between successive neurons. The peak shift across neurons was simulated as a linear function. Simulated data with linear shifts resulted in covariance matrices with a thick main diagonal pattern (Supplementary Fig. 3).

### Posterior score computation

To generate a continuous, trial-by-trial estimate of the information conveyed about task factors in population activity, we measured classifier posterior probability. In the saccade-distraction task, there were two factors related to items stored in working memory, object location and object identity. We fit separate logistic regression classifiers with lasso regularization (penalty coefficient lambda = 0.001) to decode each of these factors—sample object location (upper vs. lower) and identity (object 1 vs. object 2)—at each time point. For each classifier fit, trials were balanced between the two decoded classes by selecting a random subset of trials from the larger class. This step was repeated five times and results were averaged across these repeats. We evaluated the classification using leave-one-out cross-validation, and for each cross-validation fold, computed the classifier posterior probability for the held-out trial. The posterior score was the non-thresholded value of the logistic function corresponding to the true class of the held-out observation. This score reflects how confident the classifier was in choosing the true class label on that trial. A score of 0.5 corresponds to chance. Scores > 0.5 correspond to choosing the correct true class label on a given trial; scores < 0.5 correspond to choosing the incorrect class.

We then related posterior probability scores to the angle of neural population rotations in the saccade-distractor. Scores were computed separately for each hemisphere, area (dlPFC, vlPFC), mid-delay saccade direction (left→right, right→left), and session, and then averaged over the last half (400 ms) of the post-saccade delay (Delay 2). For each of these computations, we discretized scores into 10 equal-length bins of length 0.1. Spiking activity from the full Delay 2 period of all trials in each posterior probability bin were submitted to jPCA analysis as described above, and used to compute an angle of rotation. Thus, each hemisphere, area, saccade direction, session, and probability bin contributed a single (posterior probability score, angle of rotation) value pair. We computed the mean rotation angle (± SEM) for each probability bin (Supplementary Fig. 10), and assessed their relationship using Pearson correlation.

### Significance testing of temporal covariance

At first, we computed the baseline covariance using the trial-averaged spike-rate data during the fixation period in the saccade-distractor task. The fixation period lasted for 700 ms (= 14 time bins, each time bin = 50 ms). Each temporal covariance matrix was 𝐶_𝑏_ 𝜖 *R*^14×14^. We discarded the covariance values along the main diagonal of 𝐶_𝑏_ and considered only the off-diagonal values, which were the covariances of pairwise time bins. We computed the average of the off-diagonal values. We pooled the average values across all the sessions and averaged them to obtain a baseline temporal covariance. Next, we computed the temporal covariance matrix using the trial-averaged spike-rate data during Delay2 (see Supplementary Fig. 8 for an example session) for individual sessions. This gave a covariance tensor 𝐶*i* 𝜖 *R*^16×16×𝑛𝑜−𝑜𝑓−𝑠𝑒𝑠𝑠*i*𝑜𝑛^ . Here, the number of time bins was 16 (Delay2 duration was 800 ms = 16*50 ms). Therefore, for each time bin, we had a vector of covariance values, 𝐶_*i*,𝑗_ 𝜖*R*^𝑛𝑜−𝑜𝑓−𝑠𝑒𝑠𝑠*i*𝑜𝑛^ ^×^ ^1^. We performed a one-sample t-test on 𝐶_*i*,𝑗_ against the null hypothesis that 𝐶_*i*,𝑗_ was not different from the baseline value.

### Traveling wave estimation

We employed an algorithm developed by Horn and Shunck to measure optical flow(Horn & Schunck, 1981; Townsend & Gong, 2018) in order to detect and characterize traveling waves from noisy multi-unit recordings. Optical flow finds the apparent motion of objects in a visual scene. In our case, the “object” of interest is the neural activity moving across the “scene” of the electrode array. The optical flow algorithm takes a pair of consecutive temporal frames and outputs a velocity vector field (flow field) that represents the direction and speed at each recording location. Here, a frame denotes the neural activity in the 2D recording array at one instant.

The wave detection algorithm occurs in three steps: 1. We computed maximum and minimum firing rates of all the neurons in the trial-averaged data in a session. The time interval *T* for wave estimation was selected. Let’s assume the selected data be *Z* 𝜖 *R*^𝑛𝑒𝑢𝑟𝑜𝑛^ ^×^ ^*T*^. The data was then normalized to range [0, 1] and scaled to [0, 255]. The data was then reshaped to the physical array dimension, which was 8 × 8 × *T* in our case. In case of missing data at an electrode in the array, we performed nearest-neighborhood interpolation to simulate the spike rate data at that electrode. 2. Using the preprocessed data, we applied the “estimateFlow.m” MATLAB function to get velocity fields (*v_x_*, *v_y_*) in consecutive instants, where 𝑣_𝑥_, 𝑣_𝑦_ 𝜖 *R*^8×8×(*T*−1)^. The direction of flow was determined by computing streamlines (“streamline.m” MATLAB function) of the flow field in each of the (*T*-1) frames. Streamlines are a family of curves whose tangent vectors form the velocity field of the flow. In short, a streamline is a path traced out by a massless particle as it moves with the flow. We used all possible starting locations (8 × 8) to generate all possible streamlines. We considered streamlines that visited at least half (4/8) of the recording electrodes across any given direction. In the *i*^th^ frame, let this set of selected streamlines be *S_i_*. We denote the number of elements in the set, *S_i_* as |*S_i_*|. For each of the streamlines in *S_i_*, we computed the vector average of velocity field vectors (*v_x_*, *v_y_*) associated with the recording electrodes it visited. This step provided a principal direction vector 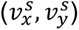of flow along each streamline 𝑠 ∈ |𝑆*i*| . The overall principal direction of flow in the *i*^th^ frame was given by the average of these across the set of all streamlines |*S_i_*|, normalized to a unit vector:

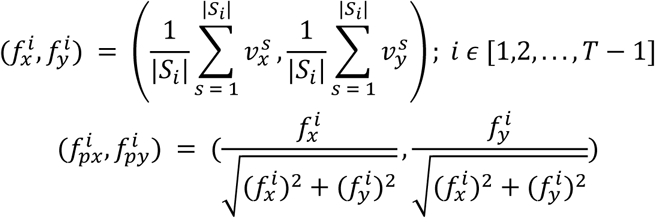

Examples of principal directions were given by black arrows in Fig. 6a.

In short, the principal direction of flow (neural activity) between two frames is the direction of the dominant flow between two instants. Note that we did not normalize the principal vectors 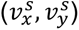 of each streamline to unit vectors. Thus, the overall principal vector 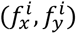 is a weighted average of per-streamline vectors 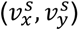 across all streamlines *s*, weighted by their magnitude. This will increase the contribution of the subset of streamlines that correspond to the dominant flow in each frame to the overall principal direction, 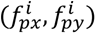.

A traveling wave has a spatio-temporal organization. The spatial organization was encoded in the principal direction. We evaluated how the principal direction was changing over time to examine the temporal organization of the wave. We computed the dot product of the normalized principal direction vectors 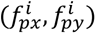 in consecutive frames to estimate the deviation of the principal flow direction over time. Traveling waves result in a relatively small deviation of the principal angle in consecutive frames, due to consistent or slowly changing temporal structure. In the absence of traveling waves, successive principal angles will be random, thus resulting in a large deviation on average of the principal angle in consecutive frames.

#### Moment to moment correlation

As mentioned earlier, spike rates were computed by binning the spike timestamps in non-overlapping 50 ms windows for analyses of state-space rotational dynamics. For analyses of traveling waves, we computed spike rates in 50 ms sliding windows in 3 ms steps, then computed the optical flow field and its principal direction vector in each temporal window (“frame”). As an estimate of the speed of flow in the *i*th frame, we computed the magnitude of its unnormalized principal direction vector: 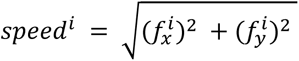. Roughly, each 50 ms time in in the state-space analysis corresponds 17 frames (17*3 ms = 51 ms) in the wave analysis. We computed the mean speed of flow of the 17 frames to obtain a speed value corresponding to each 50 ms time bin. This resulted in corresponding estimates of rotation angular velocity and wave speed at each time bin. There were 16 time bins (16 X 50 ms = 800 ms during Delay 2).

We considered a running window of length 3 (3 X 50 ms = 150 ms) to compute the correlation between the angular velocity and wave speed. We averaged across all the sessions, all working memory conditions, all hemispheres, and all animals.

#### Alternative method for traveling wave detection: Correlation of activation latency with distance

To quantify the presence of traveling waves in a way that does not depend on optical flow computations, we related the difference in activation times between pairs of neurons with their physical distance across the cortical surface (Patel et al., 2012). Any traveling wave should show an overall positive trend between pairwise activation time difference and distance. In contrast, simultaneous or spatiotemporally random activation should not exhibit such a relationship. For each neuron in the saccade-distractor task, we computed its latency to peak firing rate following the mid-delay saccade cue. An example spatial map of peak latencies from one array in a single session is shown in Supplementary Fig. 13a. For all pairs of simultaneously recorded neurons within a given array, we computed the absolute difference in their peak activation times (in ms), and their linear distance within the array (in mm). We plotted pairwise (distance, peak latency difference) for the same array and session as a scatterplot in Supplementary Fig. 13b. Finally, for each array and session, the Pearson correlation was computed between inter-neuronal distance and peak latency difference (Supplementary Fig. 13c).

### Statistics and hypothesis testing

All statistics are presented as the mean ± the standard error of the mean. In all hypothesis tests, we treated each session as an observation. We used paired t-tests (session-wise paired) in the majority of our analysis. For observations with unequal variance, we used t-test with Welch’s correction. To compare more than two variables, we used one-way ANOVA in which an F-statistic was computed. All tests were subjected to Bonferroni’s correction for multiple comparisons.

## Material availability

This study did not generate new unique reagents.

## Experimental model and subject details

The nonhuman primate subjects in our experiments were two adult rhesus macaques (*Macaca mulatta*). All procedures followed the guidelines of the Massachusetts Institute of Technology Committee on Animal Care and the National Institutes of Health.

## Data availability

The data that support the findings of this study are available from the corresponding author (E.K.M.) upon reasonable request.

## Code availability

Data analysis was performed using MATLAB R2021b (Mathworks Inc., Natick, MA). Key built-in MATLAB functions, such as estimateFlow.m and streamline.m were used for wave estimation. External toolboxes used include: jPCA (https://churchland.zuckermaninstitute.columbia.edu/ <u>content/code</u>), CircStat version 1.21.0.0 (https://www.mathworks.com/matlabcentral/ <u>fileexchange/10676-circular-statistics-toolbox-directional-statistics</u>), Fit_ellipse (https://www.mathworks.com/matlabcentral/fileexchange/3215-fit_ellipse), and TME (https://github.com/gamaleldin/TME) and CFR (https://github.com/gamaleldin/CFR) algorithms for shuffled surrogate data.

## Acknowledgements

We thank Alex J. Major, Matthew Broschard, Indie C. Garwood, Jefferson Roy, Zhen Chen, Adam Eisen, Alex Bardon, Huidi Li, and Pablo Wentz for their helpful suggestions.

## Author contributions

J.A.D., S.L.B., M.L. and E.K.M. designed the experiments. J.A.D. and M.K.M. performed the experiments and recorded the data. S.L.B. curated the data. T.B. and S.L.B. conceived and performed the analysis. E.K.M. supervised the project. T.B., S.L.B. and E.K.M. wrote the manuscript.

---

**Supplementary Figure 1.**
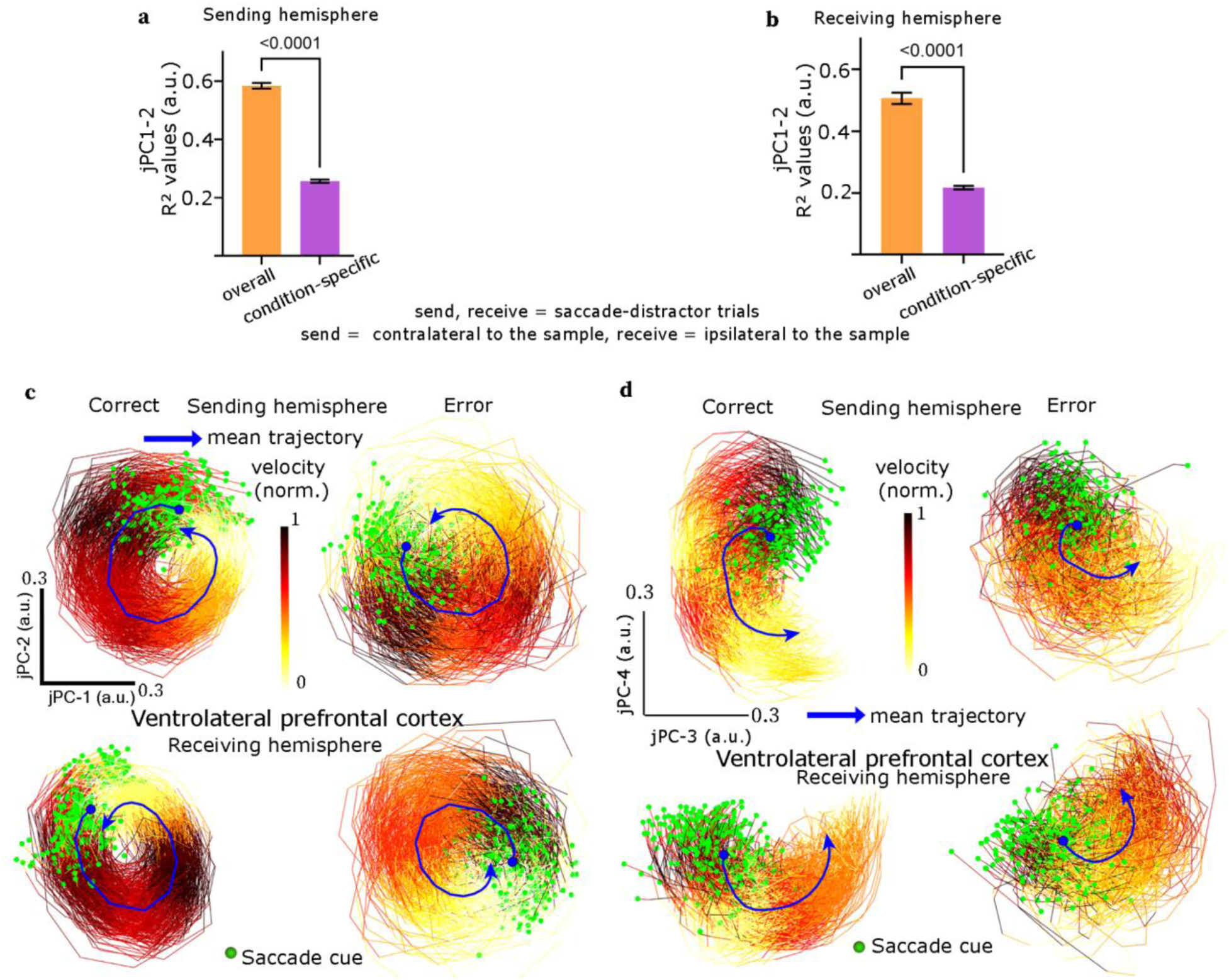
Rotational dynamics are consistent between the vlPFC and dlPFC, and stronger for raw spiking activity without mean removal (a–b) R^2^ estimates of fits during Delay 2 in the full raw spiking data (“overall”), and in spiking data with the across-trial mean activity removed, leaving only activity specific to each working memory condition (“condition-specific”). (c) Trajectories for the top two jPCs in the ventrolateral prefrontal cortex (vlPFC; compare to dlPFC results in main text Fig. 2a), for correct (left) and error (right) trials, and for the “sending” (top) and “receiving” (bottom) hemispheres. Individual trajectories start from green circles at the mid-delay saccade cue and span the entire Delay 2 (800 ms). The dark blue line indicates the mean trajectory (blue circle: saccade cue). On correct trials, the trajectories completed 360° of rotation, and then spiraled to the center (sending hemisphere = 375.03 ± 7.56°, receiving hemisphere = 363.31 ± 6.74°). On error trials, the rotations failed to complete 360° (sending hemisphere = 341.15 ± 5.13°, receiving hemisphere = 348.75 ± 8.16°). Thus, the result that the trajectories completed a fuller rotation on correct trials than error trials also holds in vlPFC (P < 0.0001, one-sided t-test; compared to dlPFC results in main text Fig. 4a). (d) Trajectories for the 3rd and 4th jPCs in the ventrolateral prefrontal cortex (vlPFC). Higher-order jPCs show weaker rotational dynamics.

**Supplementary Figure 2.**
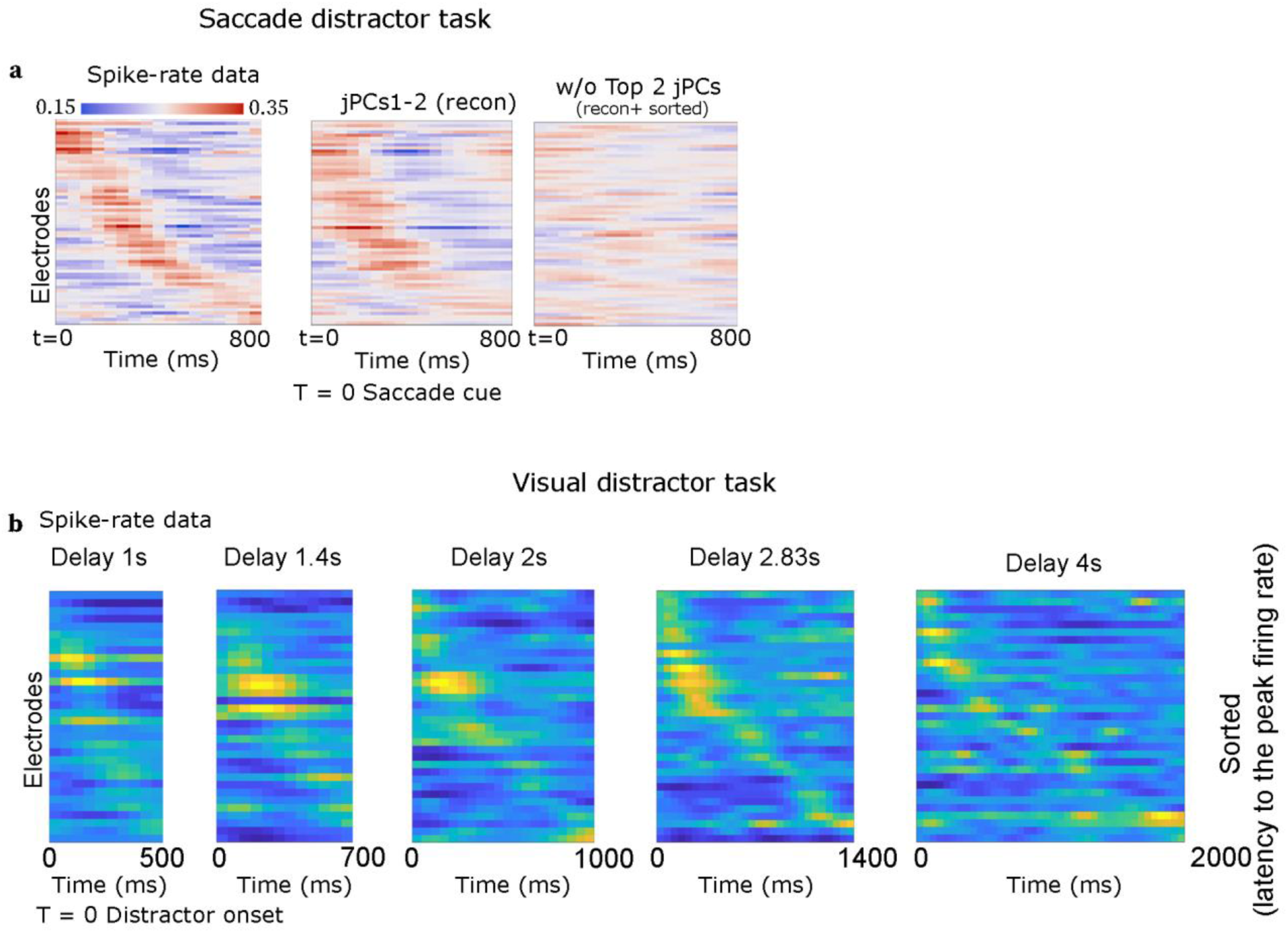
Temporal sequences of neural activation were observed in both tasks, and were well-captured by the first two jPCA components. (a) Actual spike rate data in one example session of saccade-distractor task sorted by latency to peak spike rate (*left*), the same data reconstructed from jPCs 1–2 (*middle*), and the same data reconstructed from all data *except* jPCs 1–2 (*right*). The reconstructed data were sorted using the same sorting order as the actual spike rate data. The temporal sequential structure was well-capture by the top two jPCs, and no obvious structure remained when they were removed. (b) Spike rate data from one example session of the visual-distractor task, plotted separately for each delay period length. Neurons were sorted based on the latency of their peak firing rates from the distractor onset for the shortest delay. The same sorting order was applied to the rest of the delay lengths. A temporal sequential pattern was observed for each delay length.

**Supplementary Figure 3.**
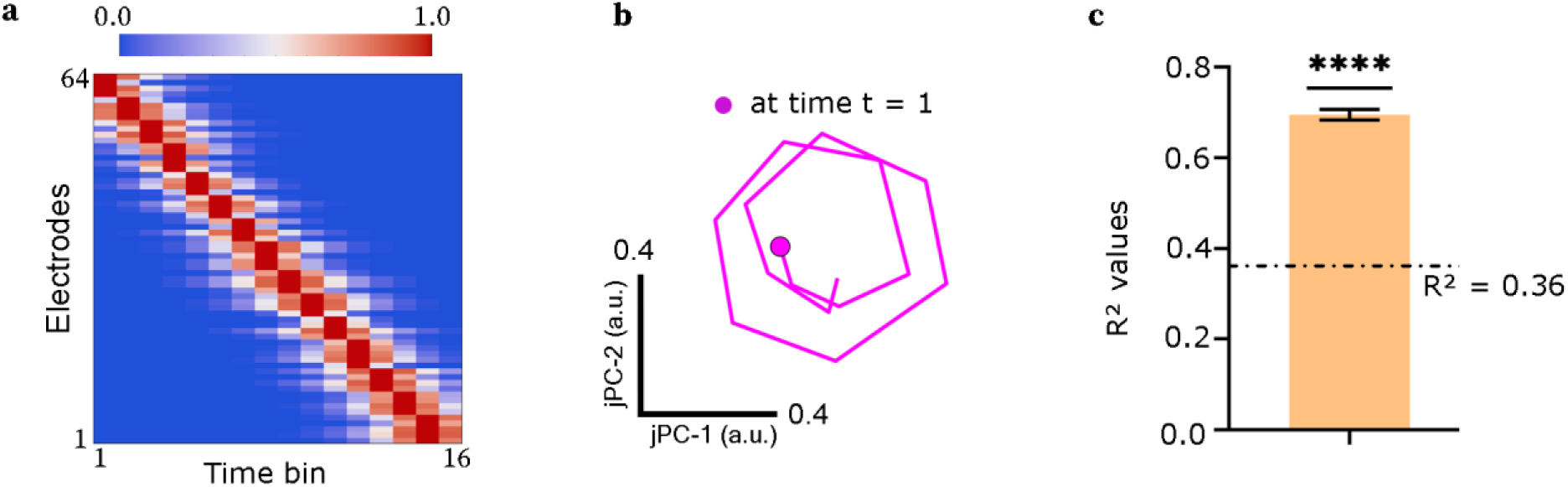
Rotational dynamics in simulated activity sequences. (a) Example of simulated sequential population activity activations (see “Sequential activation simulation” in Methods for details). The plot shows activation for a simulated population of 64 neurons (*y*-axis) across a simulated 800 ms period. Neurons are sorted by peak activation times. (b) jPCA trajectories (projections onto jPCs 1–2) corresponding to the population activity sequence. (c) R^2^ values reflecting the proportion of variance in data dynamics explained by jPCs 1–2. The simulated populations showed rotational dynamics stronger than expected by chance (dashed line), indicating that sequential activity is sufficient to induce measurable rotational dynamics.

**Supplementary Figure 4.**
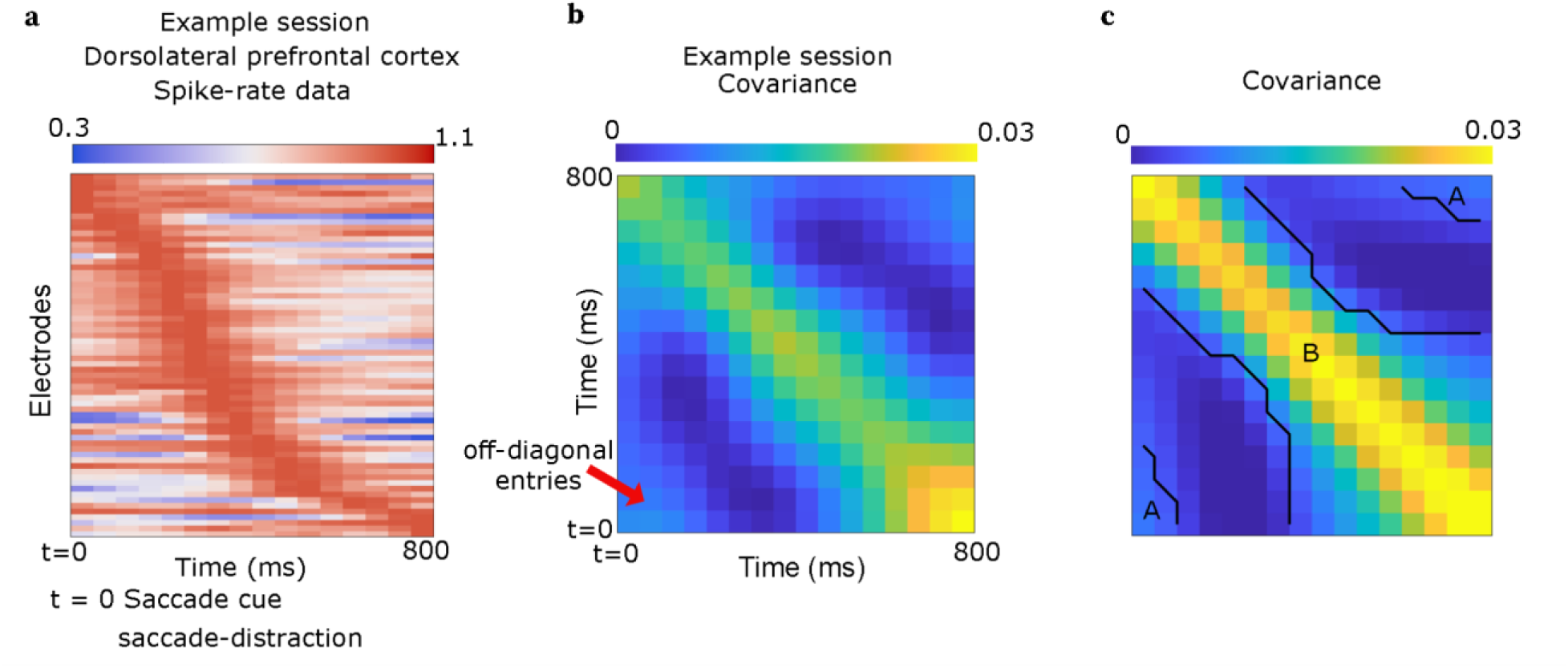
Oscillatory dynamics. (a) Spike rate during Delay 2 in right dlPFC of an example session (NHP M) of saccade-distractor task. Neurons were sorted by latency to peak spike rate. (b) Temporal covariance matrix of data in (a). In addition to the expected thick stripe along the main-diagonal (Shinn, 2023), there were off-diagonal components (red arrow), which indicate non-zero correlation between the early and late portions of the post-distraction delay. (c) Mean temporal covariance across sessions and arrays. The regions, marked by A and B, had non-zero covariance values significantly different from the baseline covariance (see Methods, “Significance testing of temporal covariance”). This provides evidence that our results are not due to either a simple non-returning sequence or to “phantom oscillations” induced by jPCA (Shinn, 2023).

**Supplementary Figure 5.**
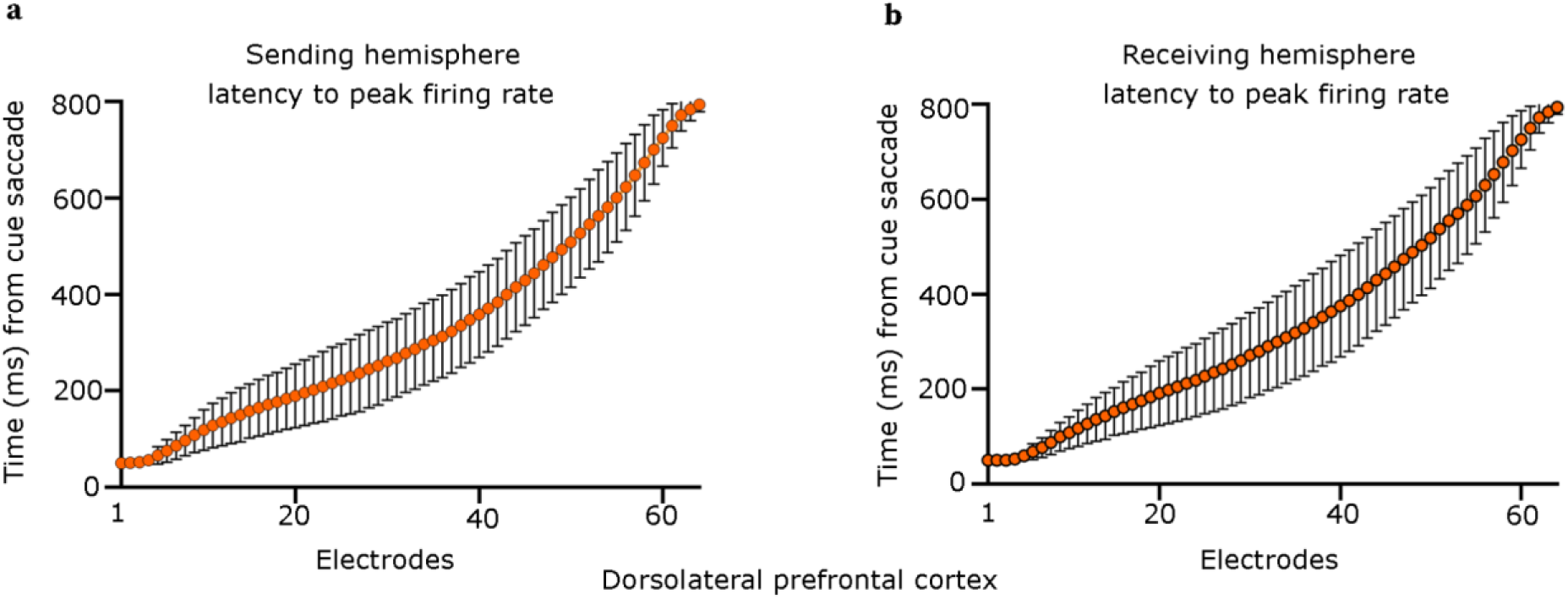
No evidence for categorical clustering of neural activity profiles. Mean ± SEM latency (n= 55 sessions) to peak firing rate after mid-delay saccade cue of each neuron from the “sending” (a) and “receiving” (b) hemisphere of dlPFC. Neurons were sorted by their peak latency independently for each session. Neurons did not show any obvious clustering into discrete groups with distinct peak activity times, but instead continuously tiled the entire delay period.

**Supplementary Figure 6.**
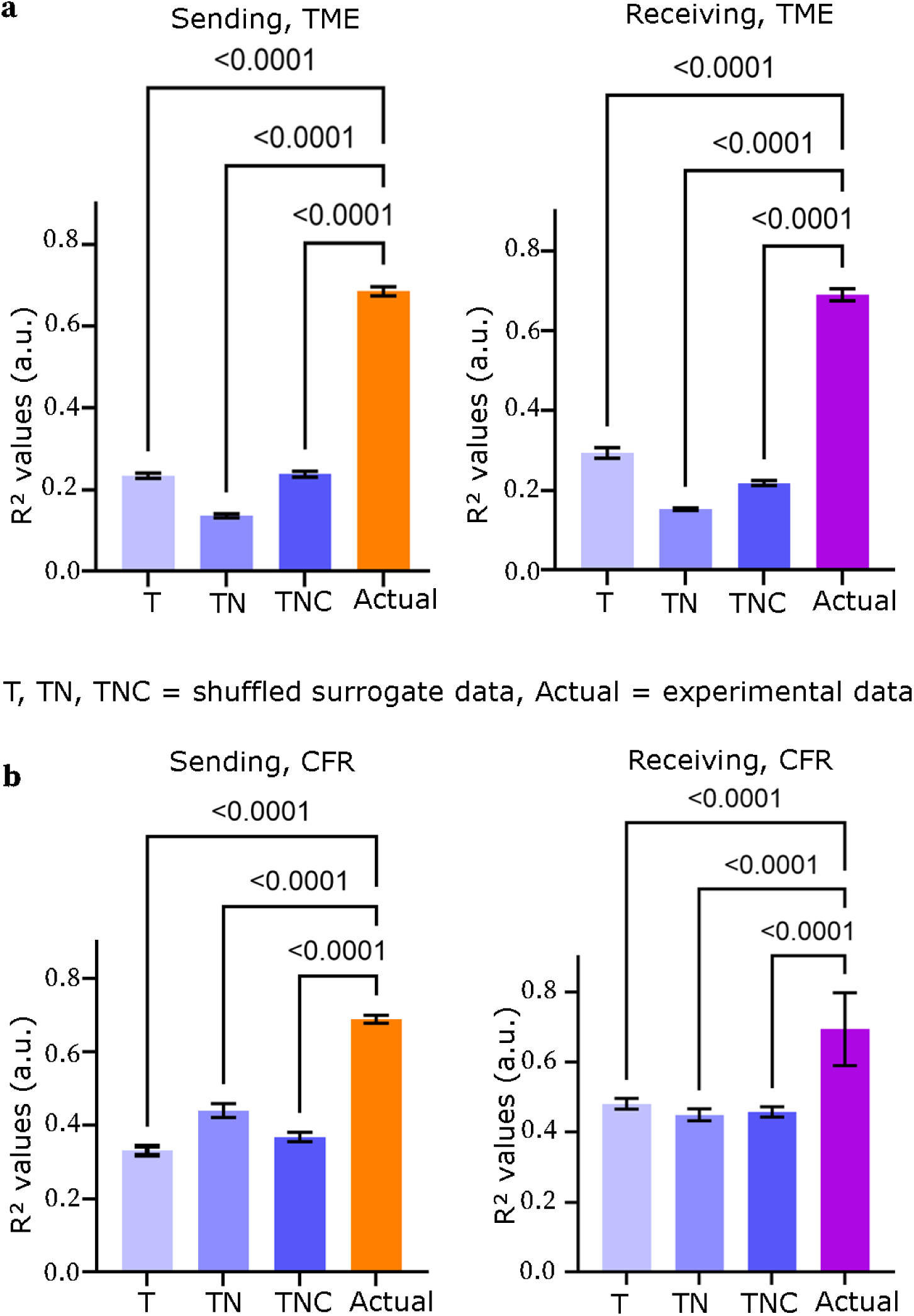
Rotational dynamics are not explained by simple features of single-neuron spiking data. (a-b) R^2^ estimates of fits of surrogate data (‘T’, ‘TN’, and ‘TNC’) and the actual data. Two algorithms, TME (a) and CFR (b), were used to generate surrogate data (dimension = 10). Here, ‘T’ = time, ‘N’ = neuron, and ‘C’ = condition. These designations refer to which dimensions of the shuffled data had their marginal mean and covariance matched to the actual data. For example ‘TNC’ means that the time, neuron, and condition dimensions were all matched to the actual data. Across all variants of this control analysis, the actual data had stronger rotational dynamics (higher R^2^) than the surrogate data.

**Supplementary Figure 7.**
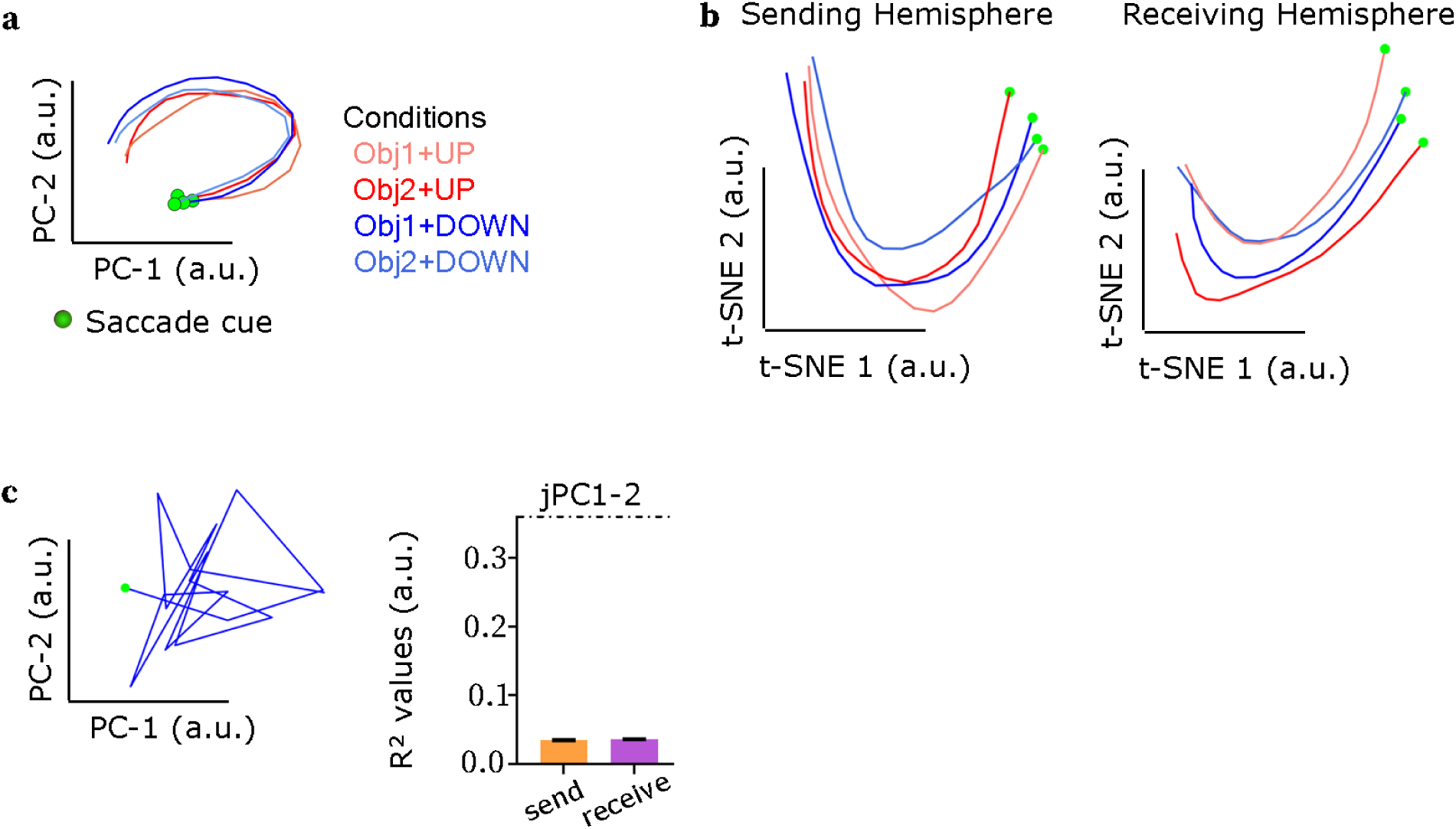
Rotational dynamics were observed with different analysis methods, but not in surrogate data or higher-order jPCA components. (a) Projections of the spike-rate data (used in (a)) on the top two PCs. Rotational structure remains visually evident, even without using methods designed to specifically capture it (jPCA). (b) Projections of the same sending (top) and receiving (bottom) hemisphere data on two t-SNE (t-distributed stochastic neighbor embedding) dimensions. (c) Projections onto the top two PCs of surrogate data that had the same PCs and variances as the actual data, but with random sampling of those PCs (see “Surrogate data generation with oscillatory PCs” in Methods) . Even though individual PCs were oscillatory, there was no rotational trajectory. This is also indicated by poor R^2^ values (bottom; compare to real data in main text Fig. 2b). Rotational dynamics additionally imply temporal orderedness in the spike rate data.

**Supplementary Figure 8.**
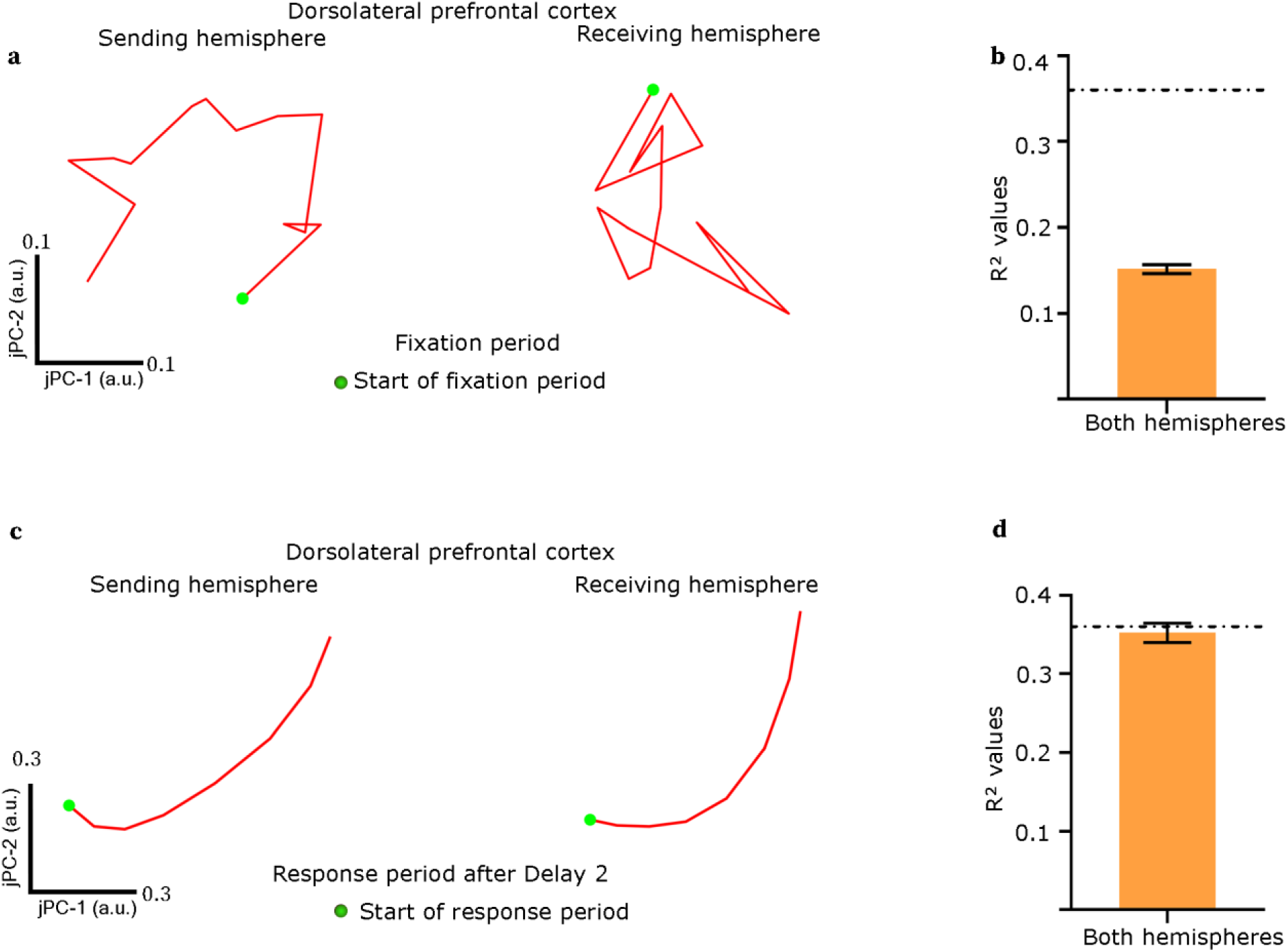
Saccades in the absence of working memory requirements do not induce rotational dynamics. We tested for rotational dynamics during the fixation and response periods, which both followed saccades but had no requirements to retain working memory across the saccade (see “Behavioral paradigm” in the Methods section). (a) Projections of fixation period spike rates from an example session and an example working memory condition onto its top two jPCs. (b) R^2^ values reflecting the strength of rotational dynamics pooled across all areas, hemispheres and sessions. (c–d) As above, for the response period. For both tested periods, there was no evidence for rotational dynamics beyond that expected by chance (dashed lines).

**Supplementary Figure 9.**
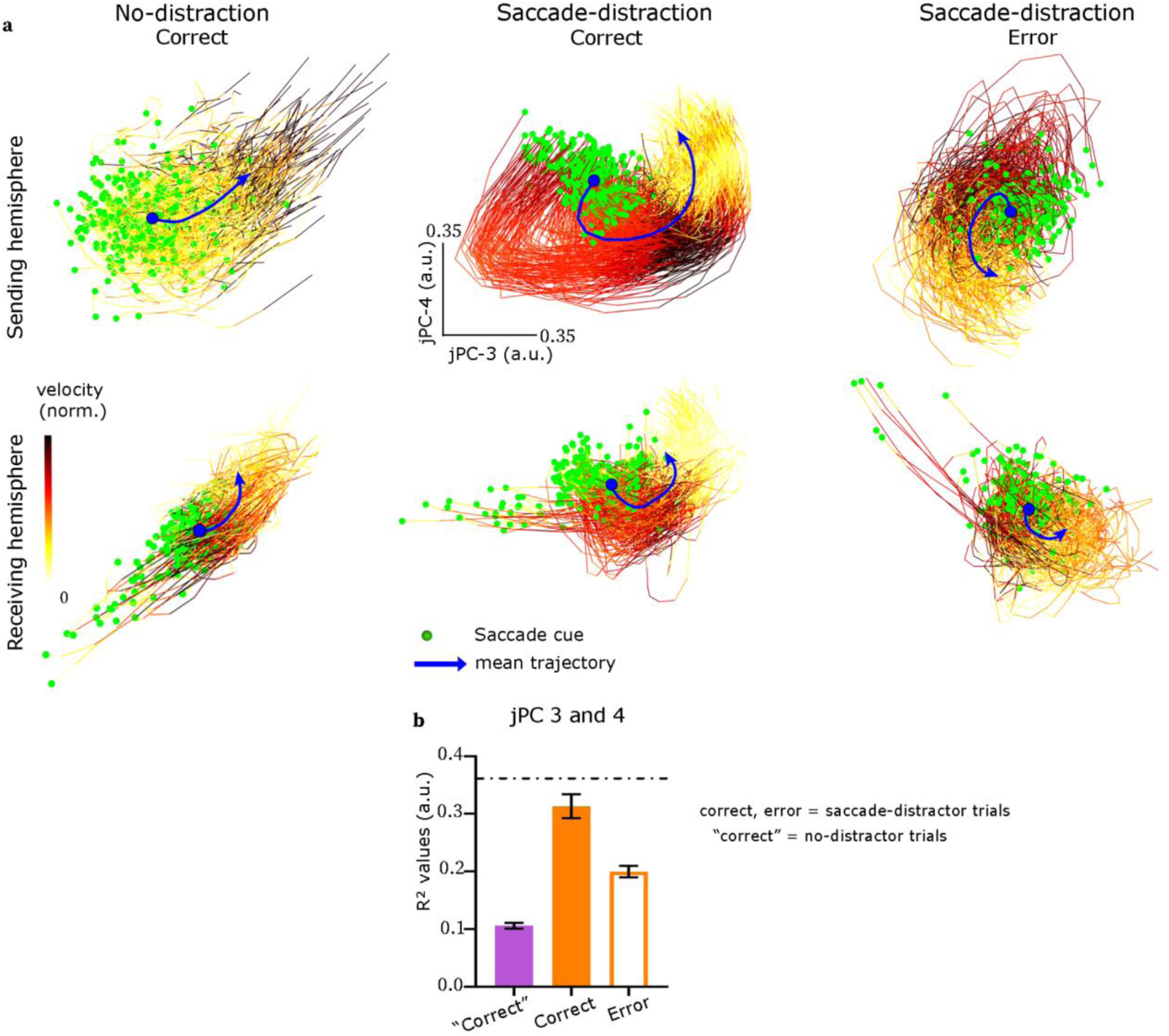
Weaker rotational dynamics in jPCs3–4 than in jPCs1–2. (a) Projections of spike rate data on the subspace of 3rd and 4th jPCs for no-distraction correct, distraction correct, and distraction error trials (compared to jPFC1–2 results in main text Fig. 2a). Individual trajectories started from green circles at the mid-delay saccade cue and spanned the entire Delay 2 (800 ms). The dark blue line indicated the mean trajectory (blue circle: saccade cue). (b) R^2^ estimates of fits for jPCs 3 and 4 (left).

**Supplementary Figure 10.**
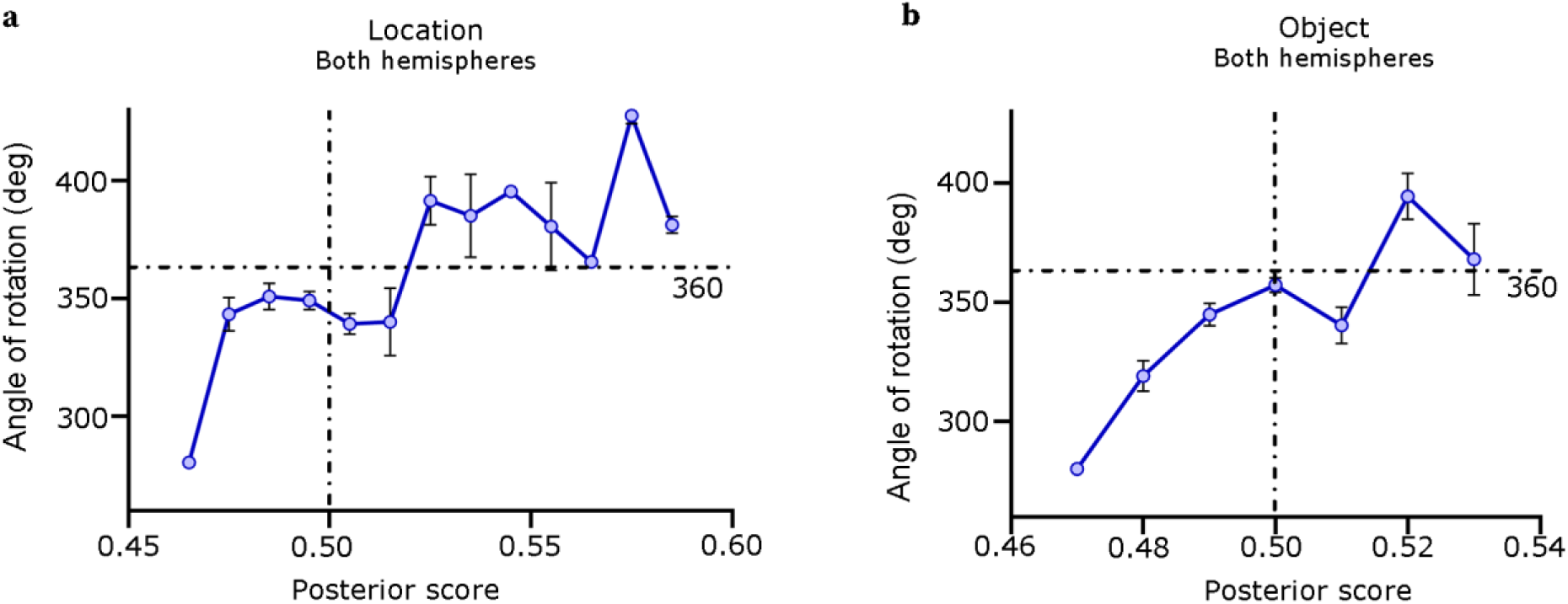
Larger rotation angles correlate with subsequent stronger population coding of items in working memory. We computed the posterior probability of classifiers trained to decode the location (a) and identity (b) of sample objects held in working memory during the final 400 ms of Delay 2. These posterior scores reflect a per-trial continuous estimate of the decoder’s confidence in choosing the actual true class. A value of 0.5 corresponds to chance (vertical dashed line). Values > 0.5 correspond to choice of the correct true class; values < 0.5 correspond to choice of the incorrect class Posterior scores were discretized into 10 equal-size bins, all trials within each bin were separately submitted to jPCA, and the total rotation angle was computed for each. Posterior scores positively correlated with rotation angles (horizontal dashed lines: 360°) for both sample object location (*r* = 0.78, *p* = 0.0014) and identity (*r* = .45, *p* = 0.012). This suggests that fuller rotations resulted in stronger population coding of information held in working memory at the end of the rotation.

**Supplementary Figure 11.**
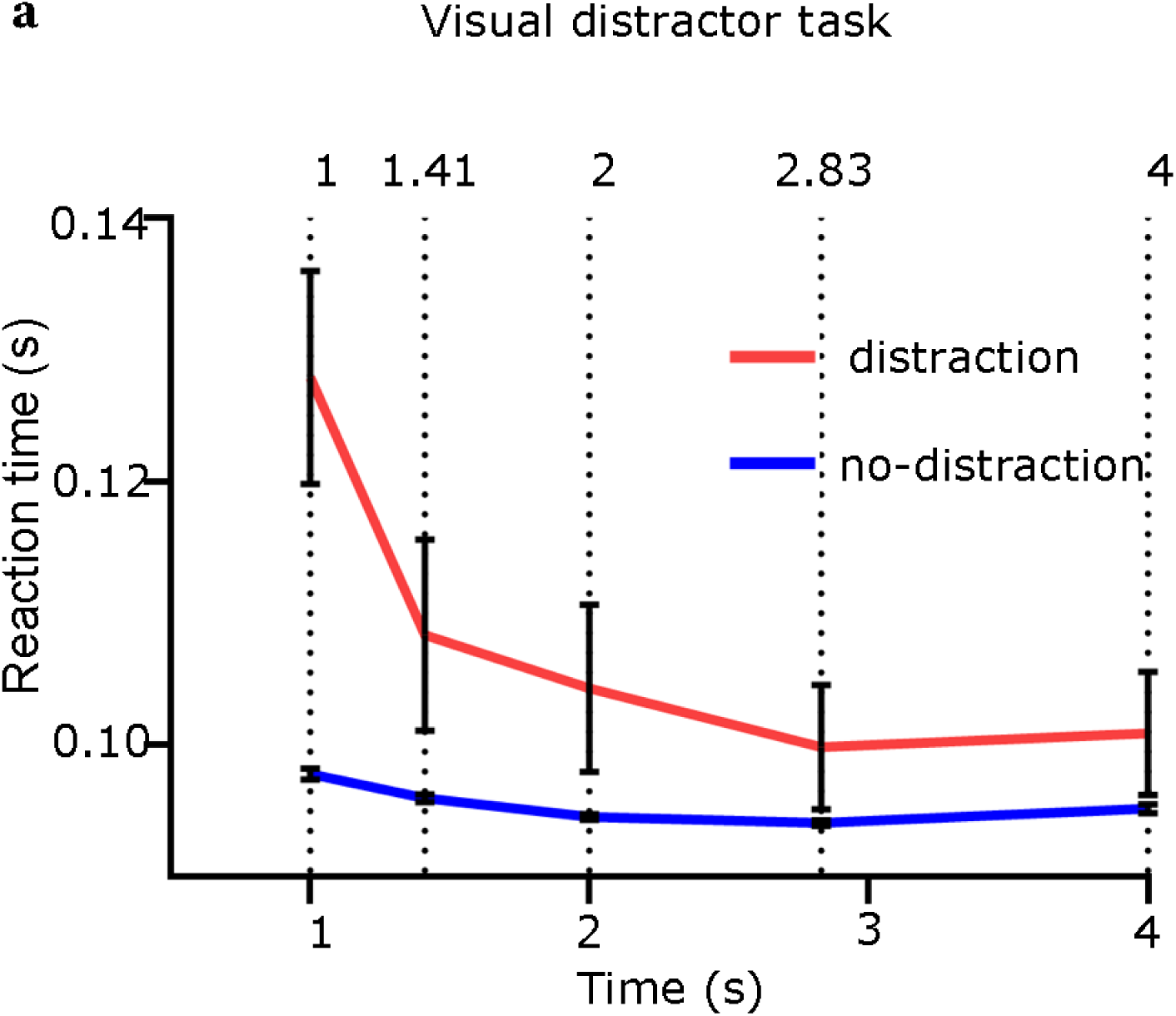
Visual distractors induced a delay-dependent behavioral impairment. Behavioral reaction time is shown as a function of working memory delay length for trials with (red) and without (blue) a mid-delay visual distractor. Trials without a distractor showed a near-constant reaction time across delay lengths. In trials with a distractor, reaction times were slowed (increased) overall, but became faster (decreased) with increasing delay length. This suggests that the distractor impaired behavior, but it gradually recovered with time.

**Supplementary Figure 12.**
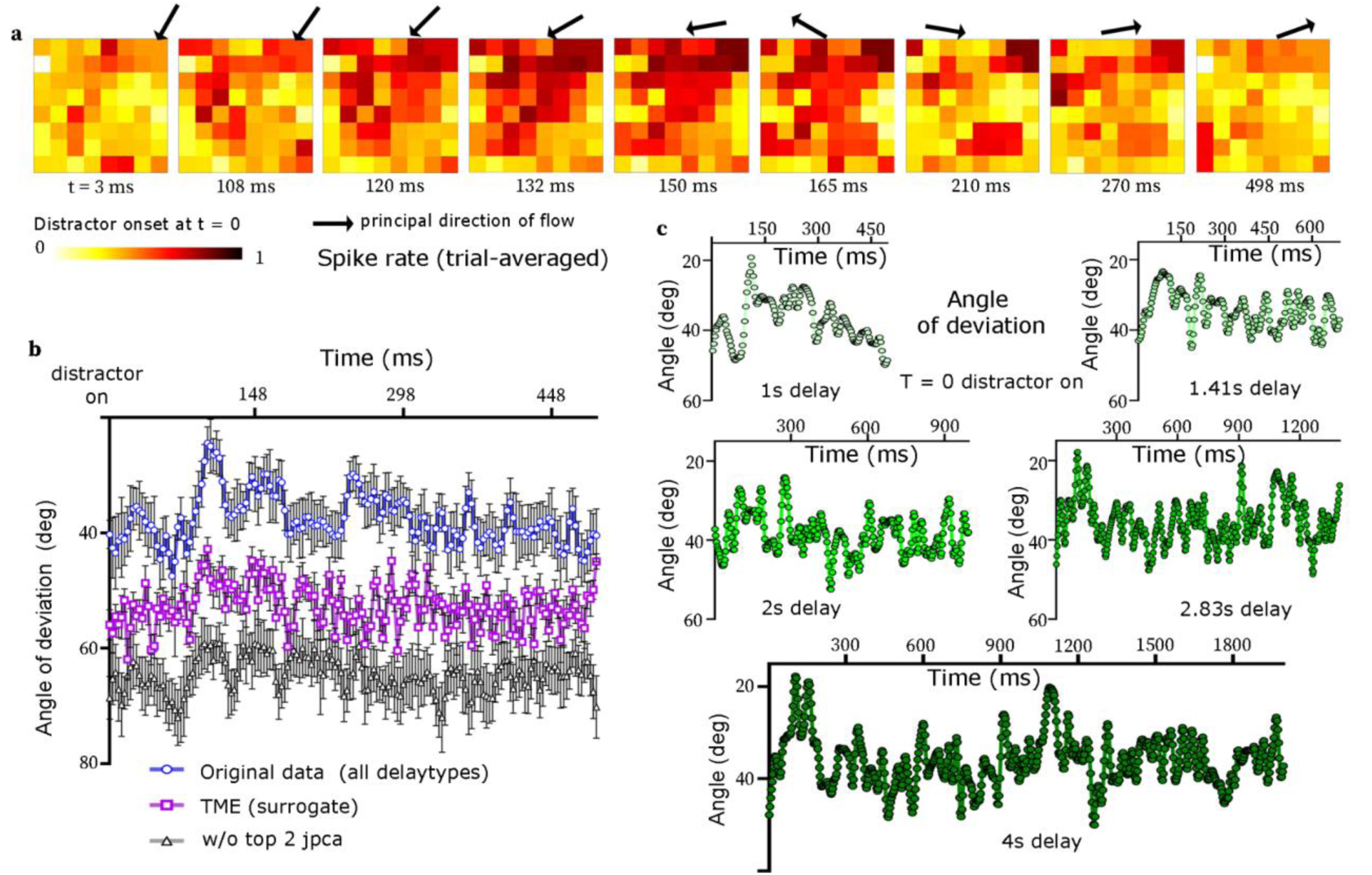
Traveling waves induced by visual distractors. (a) A wave in population-level trial-averaged spike rate caused by a visual distractor (the example was taken from the 1s delay length, and the distractor appeared mid-delay). The principal direction of flow was indicated by black arrows. (b) Correspondence between the rotations in the jPC plane and the traveling waves. Mean angular deviation of the principal direction of spike traveling waves across sessions (blue line). The same data is shown after subtracting the top two jPCs (black line) and after shuffling the data by TME algorithm (magenta line). The visual distractor also seems to induce traveling wave structure, which is captured by the top two jPCs and eliminated by data shuffling. (c) The changes in the principal direction of flow for each delay length.

**Supplementary Figure 13.**
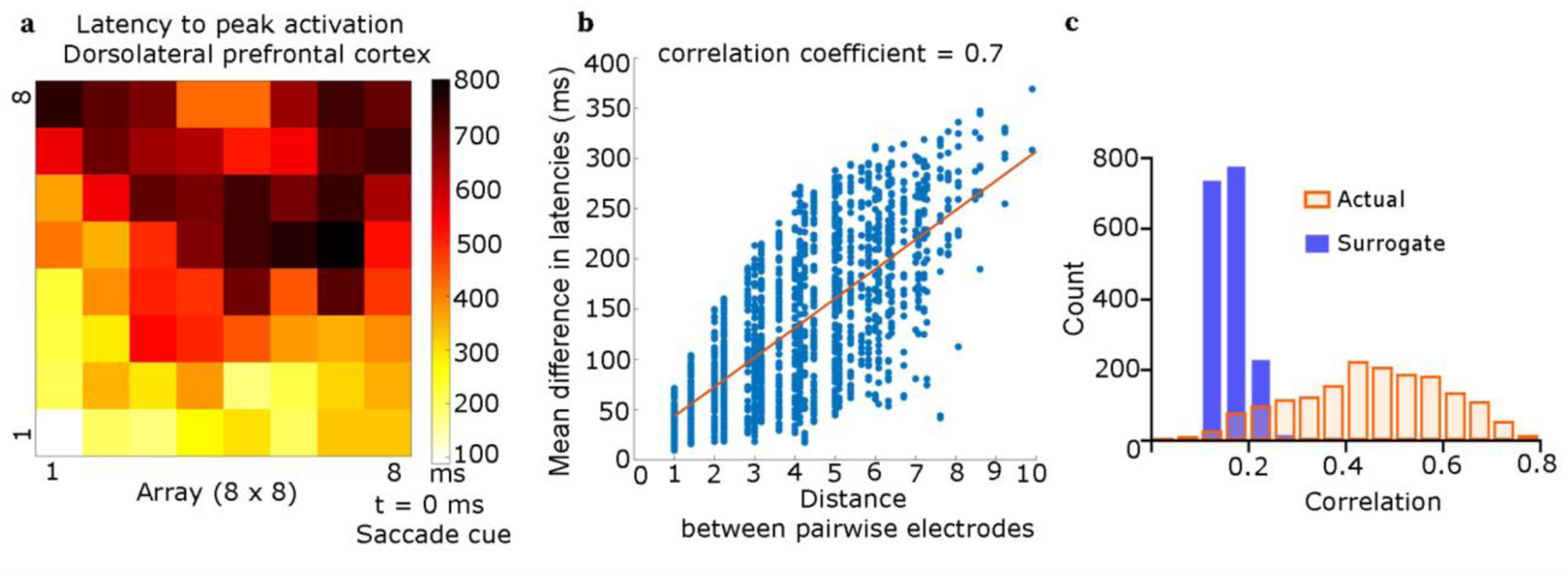
Sequential firing of neurons across space in saccade-distractor task. (**a**) Latencies to peak activation from the mid-delay saccade cue of neurons across an example 8 X 8 array in one session. In this example, there was a gradient of latency values from the bottom-left to the top-right direction, suggesting an ordered spatiotemporal propagation of spiking activity across the array. (b) Absolute differences of latency vs. spatial distance between all pairs of locations for the example data in (a). Pairwise distance significantly correlated with latency difference (*r* = 0.7, *p* = 0), consistent with a traveling wave. (b) Distribution of correlation coefficients for the actual and surrogate data across all sessions, hemispheres, and NHPs.

